# Regulation of NMDA receptor trafficking and gating by activity-dependent CaMKIIα phosphorylation of the GluN2A subunit

**DOI:** 10.1101/2021.01.11.425709

**Authors:** Xuan Ling Hilary Yong, Lingrui Zhang, Liming Yang, Xiumin Chen, Xiaojun Yu, Mintu Chandra, Emma Livingstone, Jing Zhi Anson Tan, Jocelyn Widagdo, Marta M. Vieira, Katherine W. Roche, Joseph W. Lynch, Angelo Keramidas, Brett M. Collins, Victor Anggono

## Abstract

NMDAR-dependent Ca^2+^ influx underpins multiple forms of synaptic plasticity. In the adult forebrain, the majority of synaptic NMDAR currents are mediated by GluN2A-containing NMDARs. These receptors are rapidly inserted into synapses during LTP; however, the underlying molecular mechanisms remain poorly understood. Here we show that GluN2A is phosphorylated at Ser-1459 by CaMKIIα in response to glycine stimulation that mimics LTP in primary neurons. Phosphorylation of Ser-1459 promotes GluN2A interaction with the SNX27-retromer complex, therefore enhancing the endosomal recycling of NMDARs. Loss of SNX27 or CaMKIIα function blocks the glycine-induced increase in GluN2A-NMDARs on the neuronal membrane. Interestingly, mutations of Ser-1459, including the rare S1459G human epilepsy variant, prolong decay times of NMDAR-mediated synaptic currents in heterosynapses by increasing the active duration of channel openings. Taken together, these findings not only identify a critical role of Ser-1459 phosphorylation in regulating the function of NMDARs, but also explain how the S1459G epilepsy variant dysregulates NMDAR function.

## Introduction

NMDA receptors (NMDARs) are ionotropic glutamate receptors that act as “coincidence detectors” of presynaptic glutamate release and postsynaptic membrane depolarization. NMDAR-mediated excitatory postsynaptic currents (EPSCs) mediate the flux of calcium (Ca^2+^) into the postsynaptic compartment, triggering downstream Ca^2+^-dependent signaling cascades that are crucial for neuronal development, synaptic and structural plasticity, learning and memory^1-4^. Pharmacological and genetic manipulations that disrupt the expression and function of NMDARs often cause impairments in synaptic plasticity and cognitive deficits in animal models. Importantly, NMDAR dysfunction has also been implicated in many human neurological disorders, including stroke, epilepsy, Alzheimer’s disease, neuropathic pain and schizophrenia^5^. Moreover, genes that encode NMDAR subunits are remarkably intolerant to mutations, which have been associated with various human neurodevelopmental and neuropsychiatric disorders such as epilepsy, autism spectrum disorders, intellectual disability and schizophrenia^6,7^.

The majority of NMDARs in the forebrain are heterotetramers composed of two obligatory GluN1 subunits and two identical (diheteromeric) or different (triheteromeric) GluN2 subunits^2,8-10^. Among the four different glutamate-binding GluN2 subunits, GluN2A and GluN2B, each of which confers NMDARs with distinct ion channel properties and intracellular trafficking pathways^9-11^, are highly expressed in the hippocampus and cortex^12^. The expression of synaptic NMDARs is regulated during development as they undergo a switch in their subunit composition from GluN2B-to GluN2A-containing receptors^13,14^. In the developing visual cortex, the switch in NMDAR subunit composition during the critical period can be rapidly driven by sensory experience^15^. The same phenomenon has also been observed following the induction of long-term potentiation (LTP) in acute hippocampal slices from young mice^16^, organotypic hippocampal slices^17,18^ and primary neuronal cultures^19,20^. Given that GluN2A-containing NMDARs have a higher channel open probability and a faster deactivation time than those containing the GluN2B subunit, such an activity-dependent switch in NMDAR subunit composition at synapses will have major implications for dendritic integration, circuit refinement and synaptic plasticity^21-24^. Despite this, the molecular mechanisms underlying the activity-dependent trafficking of GluN2A-containing NMDARs during synaptic plasticity remain poorly understood.

The precise subcellular localization, membrane trafficking and synaptic targeting of GluN2-containing NMDARs are largely determined by protein-protein interactions and post-translational modifications in the cytoplasmic C-terminal tails^10,25^. Sorting nexin 27 (SNX27) is a highly conserved regulator of cargo retrieval from endosomes to the plasma membrane that directly interacts with various GluN2 subunits of NMDARs through its N-terminal PDZ (postsynaptic density 95/disc-large/zona occluden-1) domain^26-28^. SNX27 forms a complex with retromer (a heterotrimer of VPS26, VPS29 and VSP35) via its direct interaction with VPS26, and acts as a cargo adaptor for retromer-mediated transport from the intracellular endosomes to the cell surface^29,30^. Genetic deletion of SNX27 causes a profound loss of total and surface NMDAR expression due to a defect in the endosomal trafficking pathway, underscoring its critical role in regulating NMDAR recycling in the brain^31^. The high affinity binding of the SNX27 PDZ domain to its cargo molecules generally involves the formation of an ‘electrostatic clamp’, formed constitutively by acidic residues at the (-3) and (-5) positions upstream of the PDZ binding motif, or alternatively by phosphorylation of serine or threonine residues in these positions^27^. The corresponding amino acid at the (-5) position within the GluN2A C-terminal tail is a serine residue (Ser-1459, see Fig. 1A), which has recently been shown to be a substrate of Ca^2+^/calmodulin-dependent kinase IIα (CaMKIIα)^28^. Importantly, a rare genetic variant that involves the S1459G substitution has been reported in a patient with epilepsy^32^. Although the phosphorylation state of Ser-1459 can regulate the interaction between GluN2A and SNX27, and receptor trafficking under basal conditions^28^, the role of Ser-1459 phosphorylation in controlling the activity-dependent endosomal recycling of NMDARs during synaptic potentiation remains unknown.

**Fig. 1.**
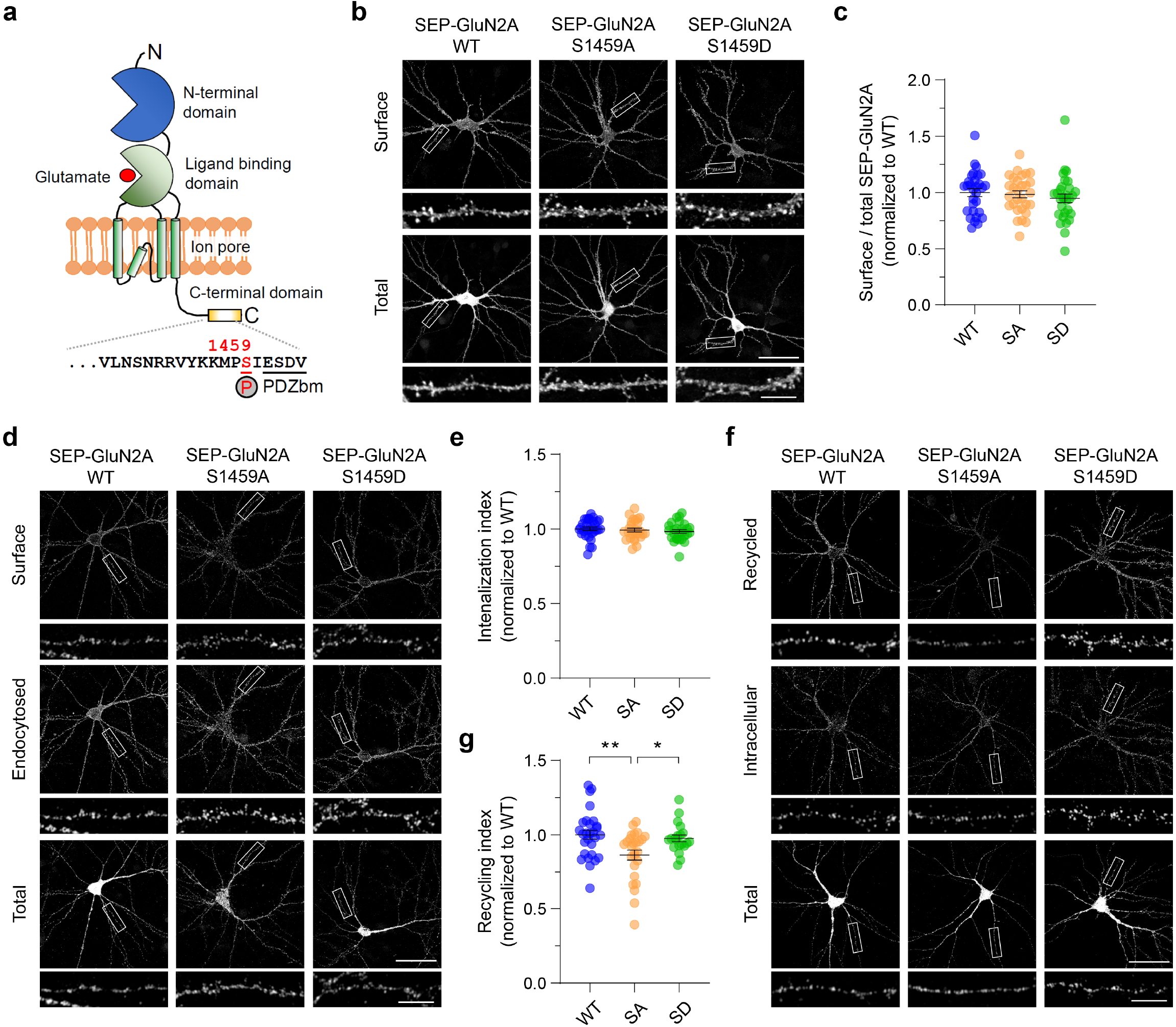
GluN2A Ser-1459 phosphorylation state regulates NMDAR recycling. **a** Schematic domain structure of a single GluN2A subunit depicting the CaMKIIα phosphorylation site, Ser-1459, which is located adjacent to the C-terminal PDZ binding motif (PDZbm). **b** Phosphorylation of Ser-1459 does not regulate GluN2A surface expression under basal conditions. Primary hippocampal neurons were transfected with plasmids encoding SEP-GluN2A, either wild-type (WT), or the phospho-deficient S1459A (SA) or phospho-mimetic S1459D (SD) mutant at DIV12. Representative images of surface and total SEP-GluN2A in a neuron from each group, together with enlarged images of the boxed regions, are shown. Scale bars, 50 μm and 10 μm (enlarged images). **c** Quantification of the surface/total GluN2A ratio normalized to the value of control neurons expressing SEP-GluN2A WT. Data are presented as mean ± SEM (WT, *n =* 30 neurons; S1459A, *n* = 29; and S1459D *n* = 30; from 3 independent cultures). **d** Phosphorylation of Ser-1459 does not affect GluN2A internalization under basal conditions. Transfected neurons were incubated with rabbit anti-GFP antibodies at 25°C for 10 min and were allowed to internalize in the medium at 37°C for 30 min. **e** Quantification of the SEP-GluN2A internalization index normalized to the value of control neurons expressing SEP-GluN2A WT. Data are presented as mean ±SEM (WT, *n =* 30 neurons; S1459A, *n* = 27; and S1459D *n* = 27; from 3 independent cultures). **f** The phospho-deficient S1459A mutant reduces the constitutive recycling of GluN2A. Transfected neurons were incubated with rabbit anti-GFP antibodies and receptors were allowed to internalize for 30 min as above. Non-internalized SEP-GluN2A was blocked with anti-rabbit Fab fragments. Neurons were transferred back to 37°C for 30 min to allow for internalized receptors to recycle back to the plasma membrane. **g** Quantification of the SEP-GluN2A recycling index normalized to the value of control neurons expressing SEP-GluN2A WT that were fixed prior to receptor recycling. Data are presented as mean ± SEM (WT, *n =* 28 neurons; S1459A, *n* = 26; and S1459D *n* = 20; from 3 independent cultures). *\* P* < 0.05, *\*\* P* < 0.01 (one-way ANOVA with Tukey’s multiple comparisons test).

In this study, we tested the hypothesis that glycine stimulation, a validated method of chemically inducing LTP in primary neuronal cultures^33^, rapidly induces CaMKIIα-mediated phosphorylation of GluN2A at Ser-1459 and consequently enhances GluN2A interaction with SNX27, which is required for upregulation of GluN2A-containing NMDARs on the neuronal plasma membrane during synaptic potentiation. We also investigated the effects of Ser-1459 phosphomutants, as well as the rare S1459G human epilepsy variant, on NMDAR-mediated EPSCs by using an engineered “artificial” synapse preparation that allows us to examine biophysical properties by NMDAR with a defined subunit composition^34^. Finally, we also performed single channel recordings from excised HEK293 membrane patches containing diheteromeric GluN2A-NMDARs to investigate the effects of Ser-1459 mutations on the gating mechanism of NMDA channels.

## Results

### The phosphorylation state of Ser-1459 regulates the basal recycling of GluN2A-containing NMDARs

Phosphorylation of NMDAR subunits in the C-terminal domain is a common mechanism controlling receptor trafficking and function^25^. A recent study has shown that GluN2A is phosphorylated on its C-terminal domain at Ser-1459 by CaMKIIα both *in vitro* and in cells^28^, a finding which we confirmed here (Supplementary Fig. 1). Co-expression of truncated CaMKIIα (tCaMKIIα) lacking the regulatory domain, which is constitutively active, and GST (glutathione S-transferase)-tagged GluN2A-C-tail (residues 1364-1464) in heterologous HEK293T cells led to a robust increase in GluN2A phosphorylation on Ser-1459 as detected by a specific antibody against GluN2A phospho-S1459 (Supplementary Fig. 1a, b). Pharmacological activation of protein kinase A (PKA) or protein kinase C (PKC) with forskolin or phorbol ester (PMA) did not affect the levels of GluN2A phosphorylation at Ser-1459 (Supplementary Fig. 1c, d). The immunoreactivity was completely abolished in cells expressing the GST-GluN2A-C-tail S1459A mutant, confirming the specificity of the phospho-antibody.

To determine the effect of Ser-1459 phosphorylation on the steady-state expression of surface GluN2A, we transfected rat primary hippocampal neurons with super ecliptic pH-sensitive GFP (SEP)-tagged GluN2A, either wild-type or the S1459A phospho-deficient or S1459D phospho-mimetic mutant, and performed a surface staining assay with anti-GFP antibodies that recognize the extracellular SEP. No significant differences in the levels of surface GluN2A were observed across genotypes (Fig. 1b, c). Next, we carried out an antibody-feeding assay with GFP antibodies to measure the degree of SEP-GluN2A internalization and recycling in live transfected hippocampal neurons over a period of 1 h (30 min internalization plus 30 min recycling). Although we did not observe any significant differences in the amount of endocytosed SEP-GluN2A among the three transfected groups (Fig. 1d, e), the GluN2A-S1459A phospho-deficient mutant caused a marked reduction in the levels of NMDAR recycling back to the plasma membrane compared to wild-type or the GluN2A-S1459D phospho-mimetic mutant (Fig. 1f, g). These results indicate that the phosphorylation of Ser-1459 promotes the endosomal recycling of GluN2A-containing NMDARs in primary hippocampal neurons under basal conditions.

### CaMKIIα-dependent phosphorylation of GluN2A at Ser-1459 enhances SNX27-retromer binding

SNX27 is known to interact with GluN2 subunits of NMDARs and play a role in receptor trafficking through the intracellular endosomal system^26-28,31^. Given that a mechanism underlying SNX27 cargo selection involves the phosphorylation of serine or threonine residues upstream of the PDZ binding motif, we predicted that phosphorylation of GluN2A at Ser-1459 would increase the binding affinity towards SNX27. To test this, we first performed an isothermal titration calorimetry (ITC) assay and measured the binding affinity of purified recombinant SNX27 PDZ domain with GluN2A peptides (Fig. 2a and Supplementary Table 1). Our results showed that the substitution of serine to glutamic acid at residue 1459 enhanced SNX27 binding (K_d_ from 46 μM to 25 μM), whereas serine to alanine substitution did not have any significant effect (K_d_ = 49 μM). As expected, a serine-to-glutamate mutation at Ser-1462 (-2 position) completely abolished the interaction between GluN2A peptide with SNX27, confirming that complex formation is dependent on the PDZ domain. We next performed a pull-down assay using total lysates of HEK293T cells co-expressing myc-SNX27 and GST-GluN2A-C-tails (residues 1213-1464) with phosphorylation deficient (S1459A) or phospho-mimetic (S1459E) mutations. Compared with wild-type GluN2A, the phospho-mimetic mutant significantly increased SNX27 binding (Fig. 2b, c). Similar results were obtained when we performed the same pull-down assay using lysates of cells co-expressing GST-SNX27 and full-length SEP-GluN2A (Supplementary Fig. 2a, b). In addition, GST-GluN2A-C-tails were able to pull-down components of the endogenous retromer complex, VPS26 and VPS35, the interaction of which was also dramatically enhanced by the GluN2A S1459E mutant (Fig. 2b, d, e). The V1464E PDZ binding motif defective mutant or GST alone failed to pull-down myc-SNX27 and the endogenous retromer complex, confirming the specificity of the assay (Fig. 2b–e). Furthermore, mutation of a single conserved histidine residue in the SNX27 PDZ binding motif interacting pocket (H112A) abolished SNX27 binding to both wild-type GluN2A and the phospho-mimetic mutant (Fig. 2f). Taken together, these data demonstrate a role for the Ser-1459 phospho-mimetic mutant in promoting the PDZ-dependent interaction between SNX27 and GluN2A.

**Fig. 2.**
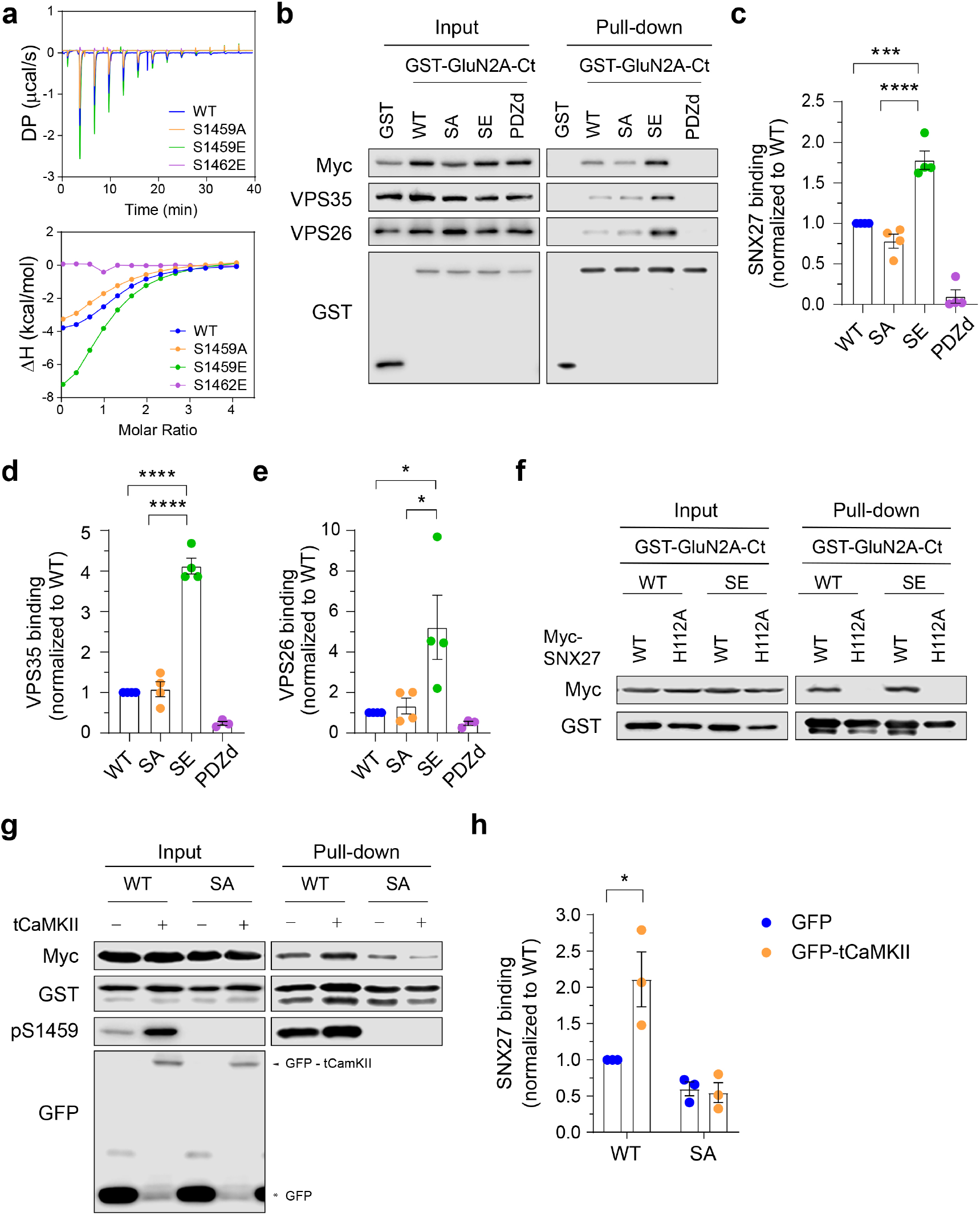
GluN2A Ser-1459 phosphorylation enhances SNX27-retromer binding. **a** ITC experiments comparing the binding of GluN2A C-terminal peptides, either wild-type (WT, blue), or the S1459A phospho-deficient (orange), S1459E phospho-mimetic (green), or S1462E PDZ binding-deficient (purple, negative control) mutant, to the purified SNX27 PDZ domain protein. The thermodynamic parameters of binding are given in Supplementary Table 1. **b** GST pull-down experiments from HEK293T cells co-transfected with plasmids encoding myc-SNX27 and GST alone or GST-GluN2A C-tails including WT, S1459A (SA), S1459E (SE) and the V1464E PDZbm mutant (PDZd). Total cell lysates (input) and bound proteins (pull-down) were resolved by SDS-PAGE, and analyzed by western blotting with specific antibodies against myc, VPS35, VPS26 and GST. **c–e** Quantification of the levels of myc-SNX27 (**c**), VPS35 (**d**) and VPS26 (**e**) binding to GST-GluN2A C-tails. Data represent mean ± SEM of band intensities normalized to WT values (*n* = 3-4; from 3 independent experiments). \**P* < 0.05, \*\*\**P* < 0.001, \*\*\*\**P* < 0.0001 using one-way ANOVA with Tukey’s multiple comparisons test. **f** GST pull-down experiments revealed that the H112A mutation in myc-SNX27 which disrupts binding to PDZbm, failed to bind both WT and the S1459E phospho-mimetic GST-GluN2A C-tails in HEK293T cells. **G** HEK293T cells were co-transfected with plasmids encoding myc-SNX27, GST-GluN2A C-tails (WT or S1459A), together with pEGFP or pEGFP-tCaMKIIα (constitutively active truncated CaMKIIα). Cells were lyzed and pulled-down with GSH-sepharose. Bound proteins and total lysates were analyzed by immunoblotting with specific antibodies against myc, GST, GluN2A pS1459 and GFP. **h** Quantification of the level of myc-SNX27 binding to GST-GluN2A C-tails. Data represent mean ± SEM of band intensities normalized to WT values (*n* = 3; from 3 independent experiments). \**P* < 0.05 using two-way ANOVA with Sidak’s multiple comparisons test. Uncropped images of the blots are shown in Supplementary Fig. 5.

To directly examine whether phosphorylation of Ser-1459 increases the binding affinity between SNX27 and GluN2A, we performed an ITC assay and found that a phosphorylated version of GluN2A C-terminal peptide bound to the PDZ domain of SNX27 with an even stronger affinity than the S1459E phospho-mimetic peptide (K_d_ = 14 μM, Supplementary Table 1). We next performed the same pull-down assay using lysates of HEK293T cells expressing myc-SNX27, GST-GluN2A-C-tails (either wild-type or S1459A phospho-deficient mutant) in the presence of GFP or GFP-tCaMKIIα. As expected, overexpression of GFP-tCaMKIIα significantly enhanced myc-SNX27 binding to GST-GluN2A-C-tail, an effect that was abolished by the GluN2A S1459A phospho-deficient mutant (Fig. 2g, h). Moreover, the same results were obtained when full-length SEP-GluN2A was used in the pull-down assay (Supplementary Fig. 2c, d). Collectively, these data provide strong evidence supporting the role of CaMKIIα-mediated phosphorylation of Ser-1459 in promoting the interaction between GluN2A and the SNX27-retromer complex.

### SNX27 is required for activity-dependent increase in surface GluN2A receptors

To assess whether SNX27 regulates the expression of surface GluN2A-containing NMDARs, we transduced primary cortical neurons with lentiviral particles expressing GFP alone, specific short hairpin RNA (shRNA) against SNX27 with GFP^35^ or myc-SNX27 cDNA. Surface biotinylation assays revealed that neither the gain-or loss-of SNX27 function affected the steady-state expression of total and surface GluN2A under basal conditions (Supplementary Fig. 3). Next, we examined the role of SNX27 in mediating the trafficking of pseudo-phosphorylated GluN2A (S1459D). We co-transfected primary hippocampal neurons with SEP-GluN2A (wild-type, S1459A or S1459D) and myc-SNX27 constructs, either wild-type or mutants that disrupt the interaction with the retromer complex (L65A) or the PDZ ligands (H112A)^29,36^. Quantification of the surface to total SEP-GluN2A ratio in the dendrites revealed a robust increase in the level of the SEP-GluN2A S1459D phospho-mimetic on the plasma membrane of neurons overexpressing myc-SNX27 wild-type (Fig. 3a, b). In contrast, expression of myc-SNX27 L65A or H112A mutants failed to promote surface expression of pseudo-phosphorylated SEP-GluN2A (Fig. 3a, b). Moreover, overexpression of myc-SNX27 (wild-type or mutants) did not result in significant changes in the levels of SEP-GluN2A wild-type or the S1459A mutant on the cell surface. Together, these results support the notion that SNX27 promotes the activity-induced endosomal recycling of phosphorylated GluN2A at Ser-1459 in a PDZ- and retromer-dependent manner.

**Fig. 3.**
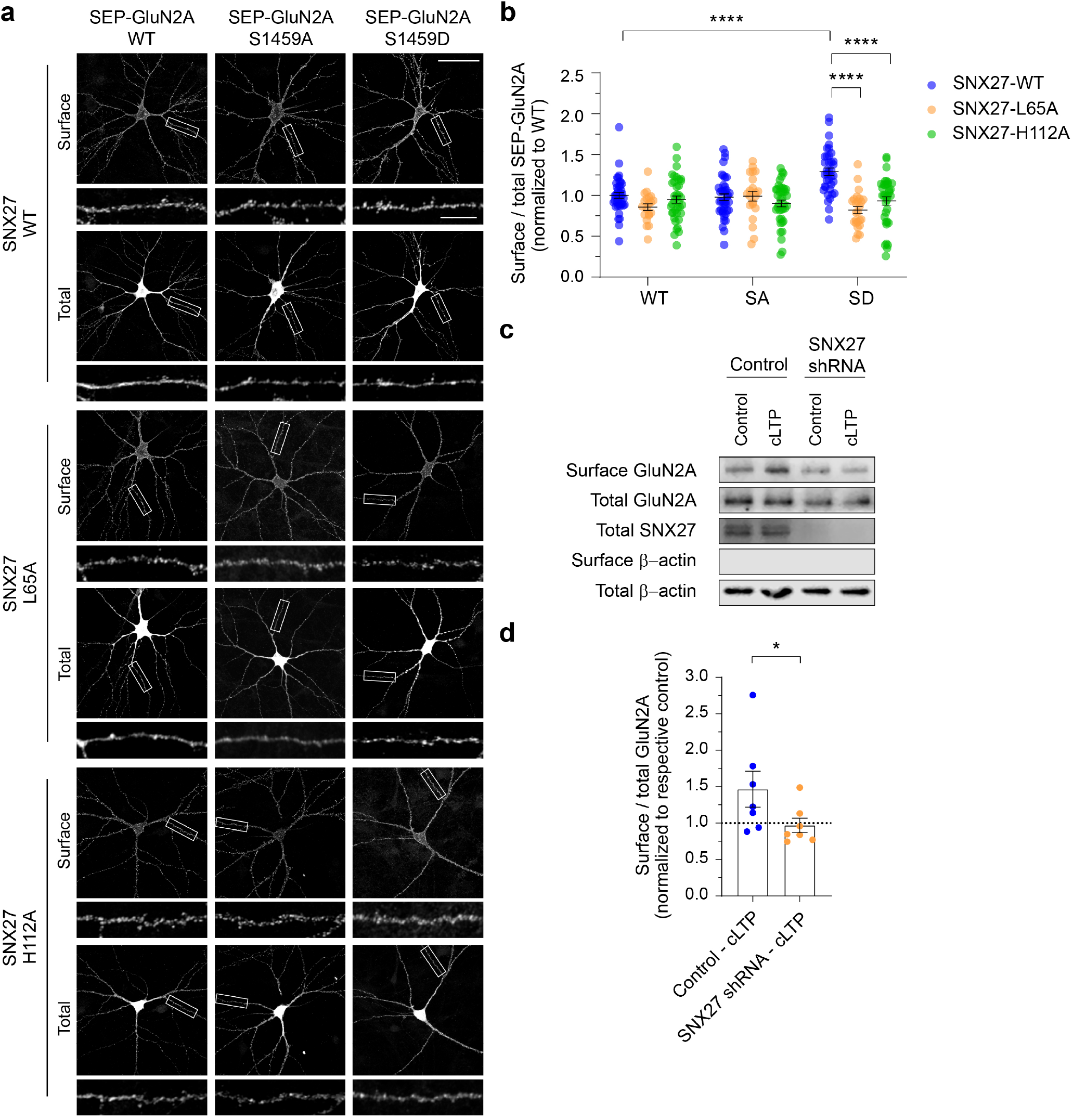
SNX27 is required for activity-dependent membrane delivery of the GluN2A-containing NMDARs. **a** At DIV12, primary hippocampal neurons were co-transfected with plasmids encoding SEP-GluN2A, either wild-type (WT), or the phospho-deficient S1459A (SA) or phospho-mimetic S1459D (SD) mutant, with various pRK5-myc-SNX27 constructs, including WT, the retromer-associated VPS26 binding-deficient mutant (L65A), and the PDZ-dead mutant (H112A). Representative images of surface and total SEP-GluN2A in a neuron from each group, together with enlarged images of the boxed regions, are shown. Scale bars, 50 μm and 10 μm (enlarged images). **b** Quantification of the surface/total GluN2A ratio normalized to the value of control neurons co-expressing SEP-GluN2A WT and myc-SNX27 WT. Data are presented as mean ± SEM (WT-WT, *n =* 39 neurons; WT-L65A, *n =* 20; WT-H112A, *n =* 38; S1459A-WT, *n* = 40; S1459A-L65A, *n* = 21; S1459A-H112A, *n* = 39; S1459D-WT, *n* = 37; S1459D-L65A, *n* = 25; and S1459D-H112A, *n* = 36; from 3 independent cultures). \*\*\*\**P* < 0.0001 using one-way ANOVA with Tukey’s multiple comparisons test. **c** Primary cortical neurons were transduced with lentiviral particles expressing GFP alone (control) or SNX27 shRNA at DIV9. At DIV15, surface biotinylation assays were performed in SNX27 knockdown and control neurons following 5 min of glycine stimulation (cLTP). The relative amounts of surface and total proteins were assessed by western blotting using specific antibodies against GluN2A, SNX27 and β-actin. Uncropped images of the blots are shown in Supplementary Fig. 5. **d** Quantification of the surface/total ratio of GluN2A in SNX27 knockdown and control neurons following glycine stimulation. Data represent mean ± SEM of band intensities relative to their respective control values (dashed line; *n* = 7; from 3 independent experiments). \**P* < 0.05 using Mann-Whitney’s test.

GluN2A-containing NMDARs are rapidly inserted into neuronal membrane during LTP^16-20^, which can be chemically mimicked (cLTP) by bath applying glycine in low Mg^2+^ artificial cerebrospinal fluid (ACSF) in primary neuronal cultures. To examine the role of SNX27 in regulating the surface targeting of GluN2A during cLTP, we performed a surface biotinylation assay and western blot on primary cortical neurons that were transduced with lentiviral particles expressing either GFP alone or SXN27 shRNA with GFP. As shown previously^19,20^, glycine stimulation resulted in a rapid and significant increase in surface GluN2A (Fig. 3c, d). However, glycine-induced enhancement of surface GluN2A was inhibited in SNX27 depleted neurons (Fig. 3c, d). These data underscore the essential role of SNX27 in controlling activity-dependent, but not basal trafficking of GluN2A-containing NMDARs in primary neurons.

### CaMKIIα-dependent Ser-1459 phosphorylation is required for glycine-induced increase in surface GluN2A receptors

Although the phosphorylation of GluN2A on Ser-1459 is dynamically regulated by neuronal activity *in vivo*^28^, it is currently unknown whether its phosphorylation by CaMKIIα can also be modulated by cLTP stimulation in primary neurons. To address this, we stimulated primary cortical neurons with glycine for 5 min and immunoprecipitated neuronal lysates with anti-GluN2A-pS1459 antibodies. Quantitative western blot analyses revealed a robust increase in the level of Ser-1459 phosphorylation following cLTP, an effect that was blocked by a pharmacological inhibitor of CaMKIIα, KN-93 (Fig. 4a, b). To corroborate these findings, we performed the same experiments in neurons that were transduced with lentiviral particles expressing Cas9-GFP alone or Cas9-GFP with a specific single guide RNA (sgRNA) against rat CaMKIIα. In agreement with the pharmacological inhibition, deleting CaMKIIα blocked glycine-induced phosphorylation of the GluN2A subunit on Ser-1459 (Fig. 4c, d). Furthermore, the application of KN-93 prevented glycine-induced up-regulation of surface GluN2A (Fig. 4e–g), suggesting that CaMKIIα-dependent phosphorylation of Ser-1459 promotes the endosomal recycling of GluN2A during synaptic potentiation. Finally, we performed a surface staining assay with anti-GFP antibodies on cultured hippocampal neurons that were transfected with SEP-GluN2A, either wild-type or the S1459A phospho-deficient mutant, following cLTP stimulation. Consistent with our surface biotinylation data, glycine treatment led to a robust increase in surface SEP-GluN2A wild-type (Fig. 4h, i). However, this effect was abolished by the S1459A mutation. In summary, these findings demonstrate a critical role of CaMKIIα-dependent GluN2A phosphorylation on Ser-1459 in promoting SNX27-retromer binding and activity-induced recycling of NMDARs during synaptic potentiation.

**Fig. 4.**
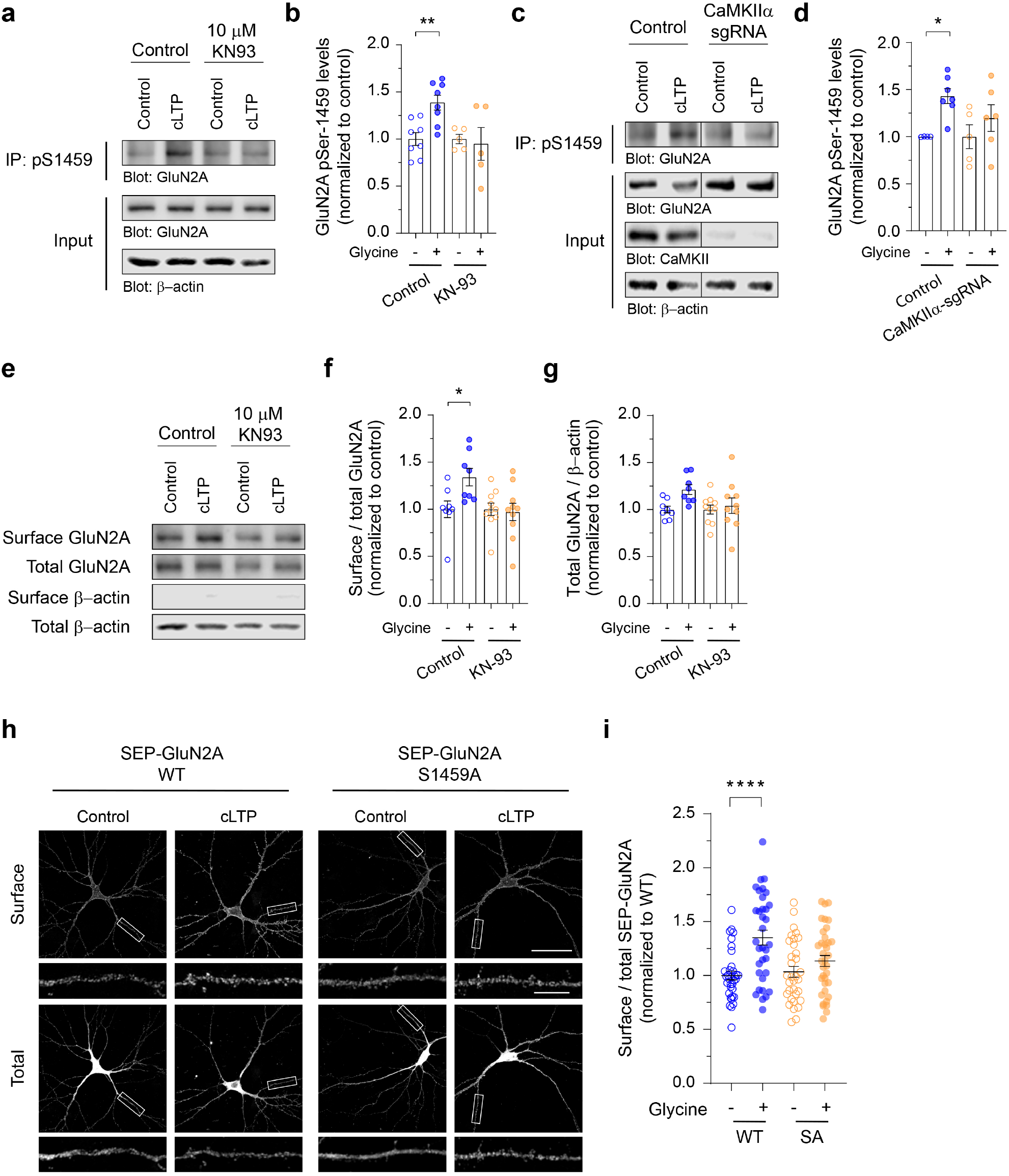
CaMKIIα-mediated phosphorylation of GluN2A at Ser-1459 is required for glycine-induced membrane delivery of NMDARs. **a** Primary cortical neurons were treated with KN-93 or water (control), and stimulated with glycine for 5 min. Endogenous GluN2A phospho-S1459 was enriched from total lysates by immunoprecipitation using anti-pS1459 antibodies, and analyzed by western blotting. **b** Quantification of the levels of Ser-1459 phosphorylation normalized to the vehicle-treated neurons (control, *n* = 8; KN-93, *n* = 5; from 4 independent experiments). Data represent mean ± SEM. \*\**P* < 0.01 using one-way ANOVA with Sidak’s multiple comparisons test. **c** Cortical neurons expressing Cas9 alone (control) or Cas9 with a specific CaMKIIα sgRNA were stimulated with glycine. The levels of GluN2A phosphorylation at Ser-1459 were determined by western blotting. **d** Quantification of the levels of Ser-1459 phosphorylation normalized to the vehicle-treated neurons (control, *n* = 4-7; CaMKIIα sgRNA, *n* = 5-6; from 3 independent experiments). Data represent mean ± SEM. \**P* < 0.05 using one-way ANOVA with Sidak’s multiple comparisons test. **e** Primary cortical neurons were treated with KN-93 or water, stimulated with glycine and subjected to surface biotinylation assays. The relative amounts of surface and total proteins were assessed by western blotting. **f, g** Quantification of the surface/total ratio of GluN2A (**f**) and the total GluN2A/β-actin ratio (**g**) in each group. Data represent mean ± SEM of band intensities relative to the control values of unstimulated vehicle-treated neurons (*n* = 8-10 per group). \**P* < 0.05 using one-way ANOVA with Sidak’s multiple comparisons test. **h** Hippocampal neurons expressing SEP-GluN2A, either wild-type (WT) or the phospho-deficient S1459A (SA) mutant, were stimulated with glycine and subjected to an antibody-feeding assay. Representative images of surface and total SEP-GluN2A in a neuron from each group are shown. Scale bars, 50 μm and 10 μm (enlarged images). **i** Quantification of the surface/total GluN2A ratio normalized to the value of unstimulated neurons expressing SEP-GluN2A WT. Data represent mean ± SEM (WT-control, *n =* 34 neurons; WT-cLTP, *n =* 34; S1459A-control, *n* = 33; and S1459A-cLTP, *n* = 36; from 3 independent cultures). \*\*\*\**P* < 0.0001 using one-way ANOVA with Tukey’s multiple comparisons test. Uncropped images of the blots are shown in Supplementary Fig. 5.

### GluN2A Ser-1459 is a Critical Residue that Regulates the Synaptic Current and Gating of Diheteromeric GluN1/2A NMDARs

In addition to its role in regulating glutamate receptor trafficking, phosphorylation of the intracellular C-terminal tail can also modulate the biophysical properties and kinetics of glutamate-gated ion channels^37^. We next evaluated the impact of Ser-1459 phosphorylation on NMDAR-mediated synaptic currents by measuring spontaneous EPSCs in HEK293 cells that form heterosynapses with mature primary cortical neurons (Fig. 5a). This experimental set up allowed us to measure EPSCs that are mediated by NMDARs with defined subunit compositions, namely diheteromeric GluN1/2A^WT^, GluN1/2A^S1459A^ or GluN1/2A^S1459D^ receptors. Our results revealed that neither the S1459A phospho-deficient nor the S1459D phospho-mimetic mutant affected the peak synaptic current amplitudes (Fig. 5a, b). However, mutations of this serine residue significantly altered NMDAR kinetics (Fig. 5a, c, d). The measurement of mean normalized currents demonstrated that these mutations slowed the 10-90% rise times (wild-type, 4.3 ± 0.4 ms; S1459A, 11.3 ± 1.1 ms; S1459D, 14.6 ± 1.5 ms, Fig. 5c). Moreover, they also slowed the decay phase of the EPSCs (wild-type, 39.1 ± 4.4 ms; S1459A, 72.7 ± 8.6 ms; S1459D, 84.5 ± 8.5 ms, Fig. 5*D*). These data clearly demonstrate that Ser-1459 is a critical determinant that regulates the kinetics of GluN2A-mediated EPSCs, regardless of its phosphorylation status.

**Fig. 5.**
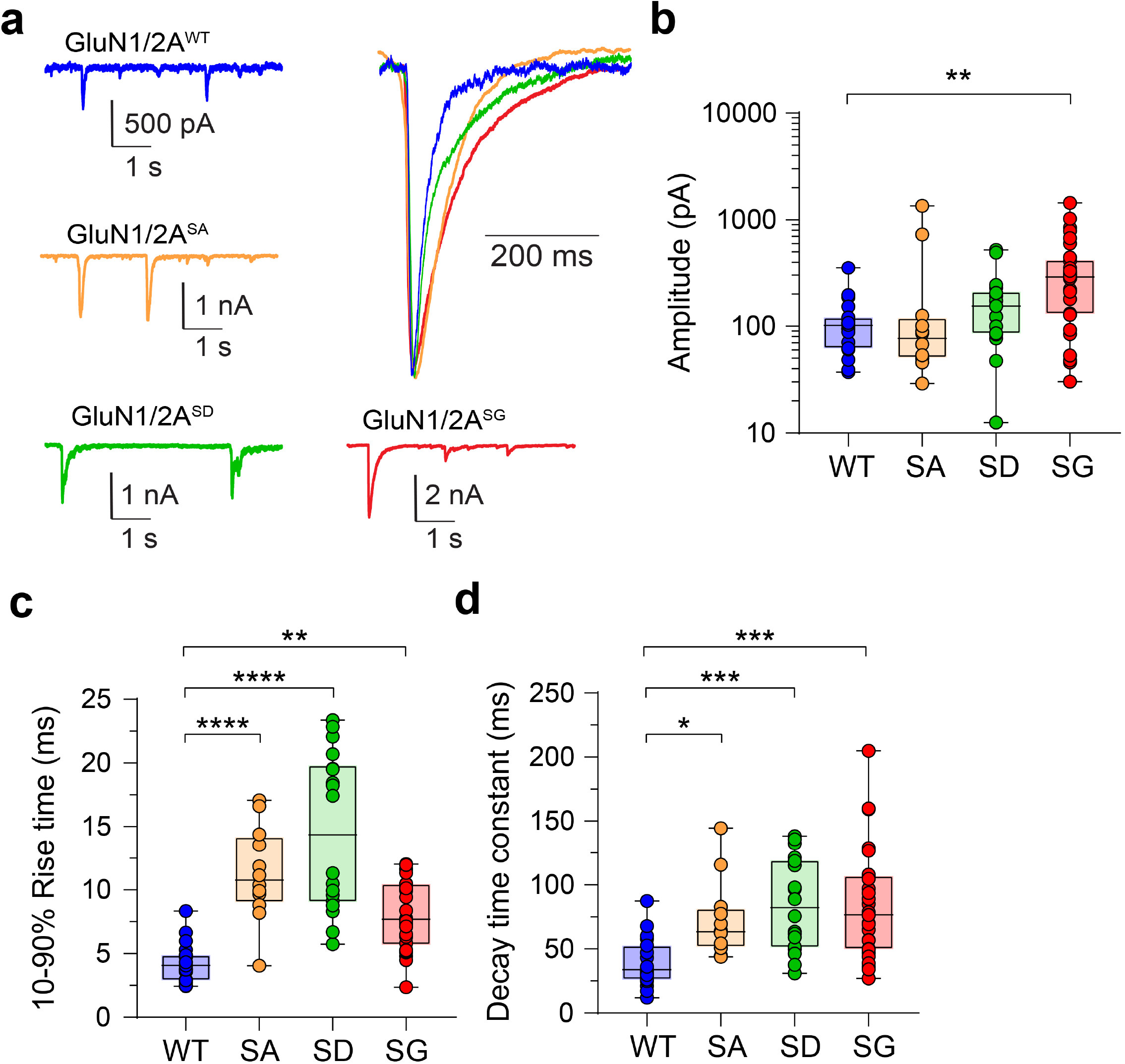
Mutations of GluN2A Ser-1459 result in prolonged excitatory postsynaptic currents (EPSCs) in heterosynapses. **a** Examples of EPSCs together with normalized, averaged current overlays (top, right) mediated by GluN1/2A^WT^ (blue), GluN1/2A^S1459A^ (orange), GluN1/2A^S1459D^ (green), and GluN1/2A^S1459G^ (red). **b–d** Summary box and whisker plots of mean synaptic current peak amplitudes (**b**), mean 10-90% activation times (**c**) and mean decay time constants (**d**) for the indicated receptors. Boxes indicate the median, 25^th^ and 75^th^ percentiles, and the whiskers mark the minimum and maximum values (WT, *n =* 19 cells; S1459A, *n =* 12; S1459D, *n* = 18; and S1459G, *n* = 30). \**P* < 0.05, \*\**P* < 0.01, \*\*\**P* < 0.001, \*\*\*\**P* < 0.0001 using one-way ANOVA with Dunnett’s multiple comparisons test.

Genetic *de novo* mutations and rare variants occurring in *GRIN2A*, which encodes the GluN2A subunit, are associated with various neurological and neurodevelopmental disorders, in particular epilepsy/seizures^6,7^. A *de novo* mutation in *GRIN2A* c.4375A>G (ClinVar, 224108), which results in the missense variant on the GluN2A Ser-1459 phosphorylation site (S1459G), has been reported in an affected individual with focal epilepsy and speech disorder^32^. We sought to determine the properties of EPSCs mediated by diheteromeric GluN1/2A^S1459G^ at heterosynapses. As expected, the S1459G mutant exhibited a slower activation time and deactivation kinetics (Fig. 5a, c, d). Surprisingly, the peak EPSCs mediated by GluN1/2A^S1459G^ were markedly larger than those mediated by GluN1/2A^WT^ (wild-type, 112.6 ± 16.8 pA; S1459G, 360.3 ± 58.5 pA, Fig. 5a, b). Taken together, these alterations to NMDAR-EPSCs would result in an increase in the charge transfer per synaptic event and represent a clear gain-of-function conferred by all three tested mutations.

To further dissect the underlying mechanisms, we examined microscopic NMDAR currents by exposing excised HEK293 membrane patches containing diheteromeric NMDARs to 1 mM glutamate and 100 μM glycine in the absence of extracellular Mg^2+^. Individual GluN1/2A^WT^, GluN1/2A^S1459G^ and GluN1/2A^S1459D^ receptors were activated in well-defined periods of variable duration (Fig. 6a). No differences in the single-receptor current amplitude were observed at −70 mV for the three receptors (wild-type, 2.60 ± 0.03 pA; S1459G, 2.68 ± 0.05 pA; S1458A, 2.69 ± 0.04 pA). Unitary current-voltage (I-V) experiments were carried out for wild-type receptors, which exhibited a mild inward rectification and a current reversal potential at −3.2 mV (Supplementary Fig. 4). Assuming that mutant receptors have the same reversal potential, we calculated the unitary conductance for all receptors and found no significant differences among groups (Supplementary Table 2). These relatively large and uniform unitary currents allowed us to measure the real-time opening and closing rates of NMDARs. Although we found no significant differences in the mean open channel probability (Po ∼ 0.65 for all groups, Fig. 6b), the open channel duration of individual active receptors was significantly enhanced in both GluN1/2A^S1459G^ and GluN1/2A^S1459D^ receptors (wild-type, 0.63 ± 0.08 s; S1459G, 1.58 ± 0.48 s; S1459D, 1.73 ± 0.23 s, Fig. 6c). Because the mean active period duration is a determinant of the ensemble current deactivation rate^38-40^, the increase in the open channel duration of mutant receptors was consistent with the observed increase in the synaptic current deactivation kinetics at heterosynapses (Fig. 5d). These results therefore suggest a clear pathomechanism: the enhanced excitatory drive caused by the increased charge transfer per synaptic event would likely promote seizure activity in the individual carrying the *GRIN2A* c.4375A>G variant.

**Fig. 6.**
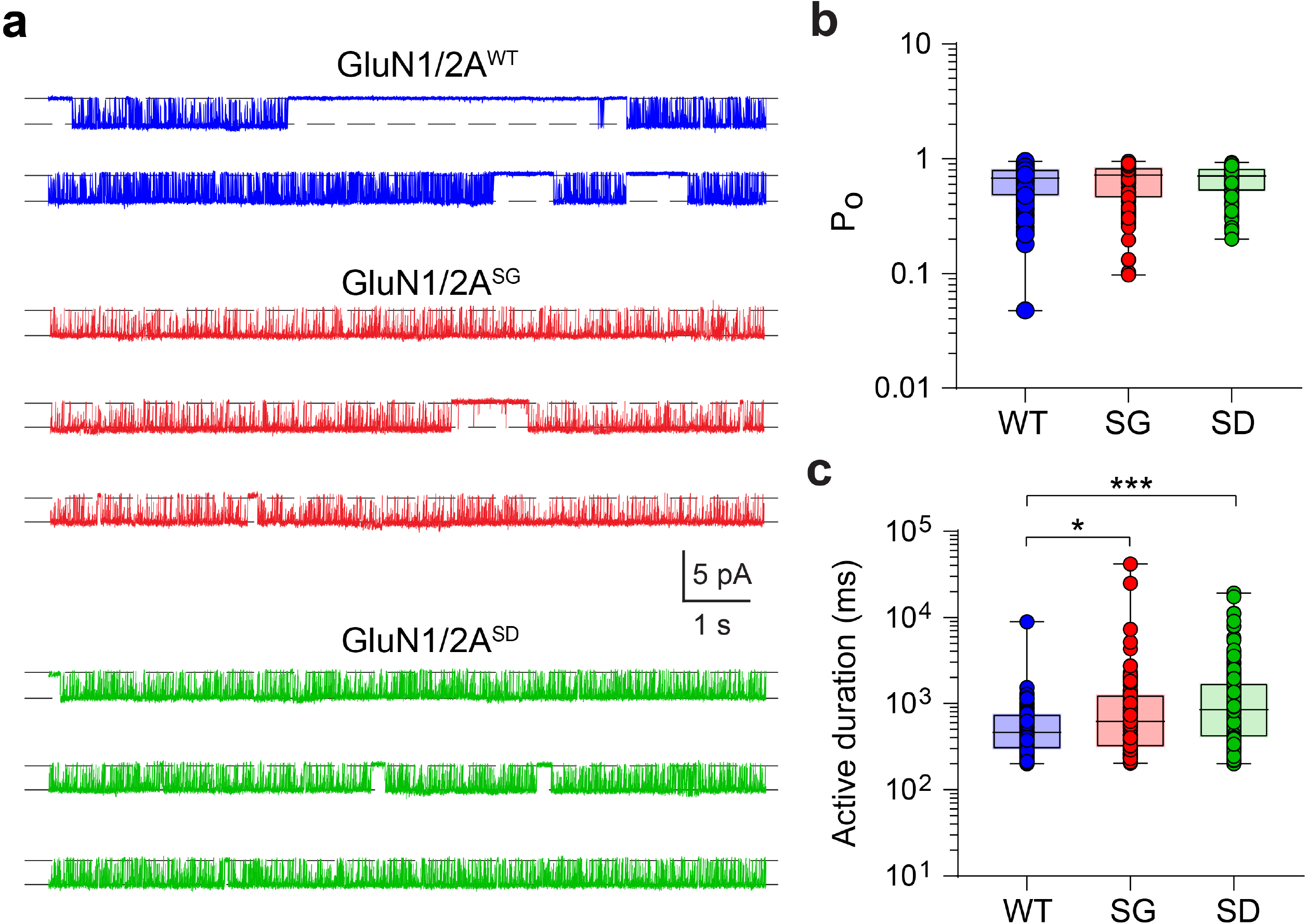
Ser-1459 mutation regulates the gating of GluN1/2A diheteromeric NMDARs. **a** Continuous sweep of single-receptor currents mediated by GluN1/2A^WT^ (top, blue), GluN1/2A^S1459G^ (middle, red) and GluN1/2A^S1459D^ (bottom, green) receptors recorded in the presence of 1 mM glutamate, 0 mM extracellular Mg^2+^ and 100 μM glycine at −70 mV. Note the similar current amplitudes, but longer active durations in the mutant receptors. **b, c** Summary box and whisker plots of intra-activation P_O_ (**b**) and active duration (**c**) of individual receptors. Boxes indicate the median, 25^th^ and 75^th^ percentiles, and the whiskers mark the minimum and maximum values (WT, *n =* 119 single receptor activations from 8 membrane patches; S1459G, *n =* 101 activations from 5 patches; S1459D, *n* = 149 activations from 16 patches). \**P* < 0.05, \*\*\**P* < 0.001 using Kruskal-Wallis one-way ANOVA with Dunn’s multiple comparisons test.

## Discussion

It is well established that NMDARs undergo a rapid switch in subunit composition from GluN2B-to GluN2A-containing receptors during LTP^16^. Despite this, our understanding of how GluN2A-NMDARs are dynamically trafficked into and out of the neuronal plasma membrane in response to neuronal activity is very limited. The present study establishes a mechanism by which activity-dependent phosphorylation of the GluN2A subunit at Ser-1459 by CaMKIIα promotes the association between NMDARs and the SNX27-retromer complex, subsequently enhancing their endosomal trafficking to the neuronal surface during cLTP (Fig. 7). In addition, we also found that molecular modifications to Ser-1459 can dramatically influence the gating of NMDARs. Because a phosphorylation-mimicking mutation (S1459D) slowed the synaptic current decay rate, we infer that the phosphorylation of Ser-1459 would do likewise and thus induce an increase in net charge transfer and Ca^2+^ influx during synaptic potentiation.

**Fig. 7.**
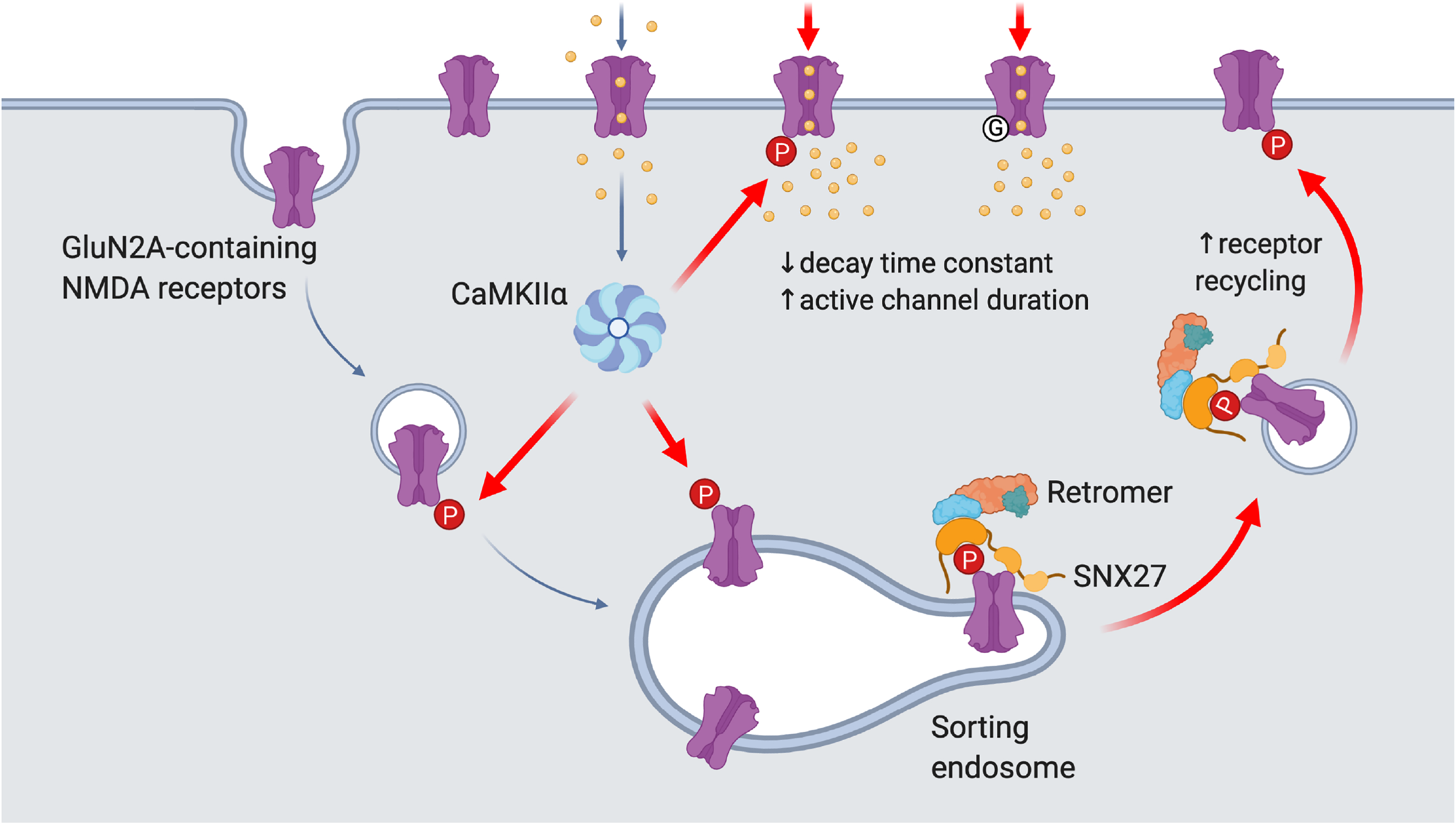
A proposed model for the role of Ser-1459 phosphorylation in regulating NMDAR functions. During LTP, NMDAR-dependent influx of Ca^2+^ into the postsynaptic compartment activates the protein kinase CaMKIIα. Phosphorylation of GluN2A at Ser-1459 by CaMKIIα in the endocytic vesicles and/or sorting endosomes promotes its interaction with the SNX27-retromer complex, thereby enhancing the rate of GluN2A recycling back to the plasma membrane. CaMKIIα can also phosphorylate existing GluN2A-containing NMDARs on the surface and affect the gating of these ion channels by prolonging the duration of single channel opening, which in turn leads to enhanced NMDAR current density and Ca^2+^ influx. Importantly, the GluN2A S1459G variant, which is associated with epilepsy, also displays augmentation in synaptic NMDAR currents due to a decrease in the receptor deactivation kinetics and an increase in active channel duration. Image was created with BioRender.com.

Unlike GluN2B, GluN2A-containing NMDARs are preferentially sorted into late endosomes for degradation^41^. The SNX27-retromer complex is one of the major endosomal retrieval complexes that prevent lysosomal degradation of many transmembrane proteins, including AMPARs^31,35,42,43^, by recycling them back to the plasma membrane^44^. Our results show that SNX27 itself is not required to maintain the steady-state expression of surface and total GluN2A in primary neurons. Overexpression of SNX27 or the GluN2A S1459D phospho-mimetic mutant does not result in increased levels of surface GluN2A, unless they are co-expressed in the same neuron. These data suggest that SNX27 and pseudo-phosphorylated GluN2A are required but not sufficient on their own to drive the endosomal recycling of NMDARs. This is in contrast with results from a recent study, which found that the GluN2A S1459A phospho-deficient mutant causes a slight but significant reduction in the surface expression of NMDARs^28^. This discrepancy could be due to several reasons, including the age and density of the neurons, experimental protocols and method of analysis. Notwithstanding these technical differences, both studies concur that phosphorylation of Ser-1459 regulates the basal recycling of GluN2A-NMDARs in neurons.

It appears that SNX27 plays a more prominent role in maintaining the steady-state expression of AMPA receptors (AMPARs) in neurons^35,45^. Although all AMPAR subunits possess C-terminal PDZ binding motifs, they do not form direct interactions with SNX27 (Refs. 27,45). Instead, their interaction is mediated by the synaptic adhesion molecule leucine-rich repeat and fibronectin type-III domain-containing protein 2 (LFRN2), which interacts with SNX27 through its PDZ binding motif^45^. LFRN2 binds to the PDZ domain of SNX27 with a low micromolar affinity due to the presence of an acidic residue at the (-3) position (Asp-786) that forms electrostatic and hydrogen bonds with Arg-58 of SNX27 (Refs. 27,45). This almost 30-fold difference in LRFN2 (K_d_ = 1.6 μM) and GluN2A (K_d_ = 46 μM) binding affinity towards SNX27 may explain the differential requirement of SNX27 in regulating the trafficking of AMPARs vs NMDARs under basal conditions.

One of the major findings of this study is that CaMKIIα-mediated phosphorylation at Ser-1459, the level of which is elevated during synaptic potentiation, enhances the interaction between GluN2A and the SNX27-retromer complex. This provides a mechanism that essentially switches the fate of internalized NMDARs from degradation to the recycling pathway in an activity-dependent manner. These results are consistent with the essential roles of SNX27 and the retromer complex in NMDAR-dependent LTP^31,42^. Therefore, our results indicate that the SNX27-retromer complex is required not only to deliver AMPARs^31,35,42,43^, but also GluN2A-containing NMDARs, to the plasma membrane during LTP.

CaMKIIα is one of the most abundant proteins in the brain and an essential regulator of NMDAR-mediated EPSCs and LTP^46-48^. Although CaMKIIα has previously been shown to interact with the distal C-terminal domain of GluN2A^49,50^, it was not until recently that a phosphorylation site (Ser-1459) was identified^28^, which we confirmed in the current study. In addition to modulating the binding of GluN2A and SNX27, the phosphorylation state of Ser-1459 also affects PSD-95 binding and the density of dendritic spines in neurons^28^. However, the impact of Ser-1459 phosphorylation on NMDAR-mediated EPSCs was not investigated. Using a HEK293–primary neuron co-culture system, we found that both S1459A and S1459D phosphomutants cause a deceleration in the synaptic current rise time and a slowing in the deactivation kinetics without affecting the amplitude of the peak currents. Mechanistically, the slower rate of receptor deactivation is due to a prolonged duration of single-channel openings. These results suggest that mutations increase the net charge transfer, establishing a role for Ser-1459 as part of the gating mechanism for GluN2A-containing receptors. It is conceivable that post-translational modification of Ser-1459 through CaMKIIα-mediated Ser-1459 phosphorylation has the ability to enhance excitatory input and Ca^2+^ signaling of target neurons during LTP. Overall, these findings further emphasize the importance of protein phosphorylation of the intracellular C-terminal domain of glutamate receptors as a general regulatory mechanism for receptor gating, which is best exemplified by the effects of GluA1 phosphorylation by PKA and CaMKIIα on AMPAR open probability and channel conductance^51-53^.

Many genetic *GRIN2A* variants have been found in patients with epilepsy. The majority of these variants are located in the extracellular and transmembrane domains where they typically result in an overall increase in NMDAR-mediated currents^6,7^. However, a small number of mutations in the intrinsically disordered intracellular C-terminal domain have also been reported, including the missense variant on the GluN2A Ser-1459 phosphorylation site (S1459G) in an individual with focal epilepsy and speech disorder^32^. The GluN2A-S1459G variant exhibits defects in SNX27 and PSD-95 binding, and consequently causes deficits in receptor trafficking, reduced dendritic spine density and a decrease in the frequency of AMPAR-mediated mini-EPSCs when overexpressed in cultured hippocampal neurons^28^. Here, we directly measured the effects of the S1459G mutation on NMDAR-mediated currents at heterosynapses. As expected, the S1459G variant altered the kinetics of receptor activation and deactivation, accompanied by an aberrant enhancement in the peak current amplitude. Single-channel recordings from excised patches confirmed the prolonged duration of single-channel opening as the underlying mechanism. Our findings therefore identify functional consequences of the S1459 variant that lead to both gain-and loss-of NMDAR function, highlighting the complex nature of a GluN2A C-terminal variant in disease etiology. Indeed, rare variants that cause mis-sense mutations in the M2 channel pore-forming loop of various NMDAR subunits have previously been reported to cause gain-of channel function accompanied by reduced surface expression of the receptors^54^. This highlights the need to study the complex effects of NMDAR rare variants on neuronal function throughout development, which will potentially be of clinical importance towards personalized medicine.

In conclusion, our findings describe dual mechanisms by which CaMKIIα-mediated phosphorylation of GluN2A at Ser-1459 regulates the trafficking and gating of NMDARs during synaptic potentiation (Fig. 7). We propose that during LTP, NMDAR-dependent influx of Ca^2+^ into the postsynaptic compartment activates the protein kinase CaMKIIα. Phosphorylation of GluN2A at Ser-1459 by CaMKIIα in the endocytic vesicles and/or sorting endosomes promotes its interaction with SNX27. The assembly of SNX27-retromer complex further enhances the binding affinity towards phosphorylated GluN2A, and subsequently increases the rate of GluN2A recycling back to the plasma membrane. CaMKIIα can also phosphorylate existing GluN2A-containing NMDARs on the surface and prolong the duration of single-channel opening, thereby augmenting the NMDAR current density and Ca^2+^ influx. On a broader scale, these mechanisms may be important in the expression and maintenance of bidirectional plasticity, dendritic integrity, learning and memory.

## Methods

### DNA constructs

The plasmid encoding SEP-GluN2A was created by subcloning rat GluN2A cDNA into the pRK5 vector and subsequently inserting SEP cDNA after the signal peptide (residues 1-34). GST-tagged GluN2A C-terminal tails were generated by amplifying GluN2A cDNA encoding residues 1213-1464 and 1364-1464 with the following primers: residues 1213-1464 FP, 5’-ATC<u>GTCGAC</u>CCATTGCAGAAGCTGCCTTTCGAAT-3’; residues 1364-1464 FP, 5’-ATC<u>GTCGAC</u>CGATAATCCTTTCCTCCACACGTAT-3’; common RP, 5’-TCGA<u>GCGGCCGC</u>TCTTAAACATCAGATTC-3’. The PCR products were subsequently cloned into the unique Sal*I* and Not*I* sites in the pCIS-GST vector. Full-length SNX27 cDNA was subcloned from pRK5-myc-SNX27 into the same pCIS-GST or FUW-myc lentiviral vector using the unique Sal*I* and Not*I* sites. GluN2A (S1459A, S1459D, S1459E, S1459G and V1464E) and SNX27 (H112A and L65A) point mutants were generated with an overlapping PCR protocol. The shRNA targeting the rat SNX27 sequence (5’-AACCAGGTAATAGCGTTTGAA-3’) was cloned into the FG12 lentiviral vector, which has been validated previously^35^. The sgRNA targeting the rat CaMKIIα sequence (5’-CTCTTCCGTGAATCGGGTGC-3’) was cloned into the pLentiCRISPR-GFP vector^48^. Other plasmids encoding untagged-GluN1^55^, HA-neuroligin-1B^56^, GFP-tCaMKIIα^57^, myc-SNX27^35^ and GST-SNX27 PDZ domain^29^ have been reported previously. The pEGFP-C1 vector was obtained from Clontech.

### Antibodies

Rabbit polyclonal antibody against GluN2A phospho-S1459 was custom-made and has been validated previously^28^. The following antibodies were purchased from commercial sources: mouse anti-β-actin (Cat# sc-47778, Santa Cruz Biotechnology), rabbit anti-CaMKII (Cat# 4436, Cell Signaling Technology), chicken anti-GFP (Cat# GFP-1020, Aves Labs); rabbit anti-GFP (Cat# 50430-2-AP, Proteintech); rabbit anti-GluN2A (Cat# 04-901, Millipore), mouse anti-GST (Cat# 66001-2-Ig, Proteintech), mouse anti-myc (Cat# sc-40, Santa Cruz Biotechnology), rabbit anti-SNX27 (Cat# 16329-1-AP, Proteintech), rabbit anti-VPS26 (Cat# ab181352, Abcam) and rabbit anti-VPS35 (Cat# 10236-1-AP, Proteintech). Polyclonal rabbit antibody against GFP (JH4030) was a gift from Dr. Richard Huganir^58^. Alexa-conjugated and HRP-conjugated secondary antibodies were purchased from Thermo Scientific and GE Healthcare, respectively.

### Isothermal titration calorimetry

Recombinant rat SNX27 PDZ domain (residues 38-135) was expressed in *E. coli* BL21(DE3) cells and purified on glutathione-Sepharose columns, as described previously^27^. ITC experiments were carried out on a Microcal iTC200 instrument in buffer consisting of 50 mM HEPES (pH 7.5), 2 mM DTT and 100 mM NaCl. Peptides (in the range of 2-50 mM) were titrated into 50-100 μM SNX27 PDZ domain at 25 °C. Data were processed with ORIGIN (OriginLab) to extract the thermodynamic parameters Δ*H, K*_d_ (1/*K*_a_) and the stoichiometry, n. Δ*G* and Δ*S* were derived from the relationship: Δ*G* = -RTln*K*_a_ and Δ*G* = Δ*H* -TΔ*S*. The thermodynamic binding parameters are reported in Supplementary Table 1.

### HEK293T cells and GST pull-down assays

HEK293T cells were grown in DMEM with 4.5g/l glucose supplemented with 10% FBS, 50 U/ml penicillin and 50 μg/ml streptomycin in a humidified 5% CO_2_ incubator at 37°C. Cells expressing functional GluN1/2A NMDARs were maintained in medium supplemented with 40 mM MgCl_2_ to avoid excessive Ca^2+^ influx and cell toxicity. GST pulldown assays were carried out as previously described^59^. Briefly, HEK293T cells were co-transfected with either pCIS empty (GST), pCIS-GluN2A-C-tails (GST-GluN2A C-tails) or pCIS-SNX27 (GST-SNX27) in combination with other plasmids by the calcium-phosphate precipitation method. To activate the PKA and PKC signaling pathways, cells were treated with forskolin (Sigma, 10 μM) or PMA (Sigma, 1 μM), respectively, for 10 min prior to cell lysis. Cells were lyzed 48 h post-transfection with ice-cold cell lysis buffer (1% Triton X-100, 1mM EDTA, 1mM EGTA, 50mM NaF, 5mM Na-pyrophosphate in PBS) supplemented with EDTA-free protease inhibitor cocktail. Lysates were centrifuged at 20,627*g* for 20 min at 4°C and incubated with glutathione agarose beads (Thermo Scientific) overnight at 4°C. Beads were washed three times with ice-cold cell lysis buffer and bound proteins were eluted with 2X SDS sample buffer at 50°C for 30 min and analyzed by western blotting. Blots were analyzed using the enhanced chemiluminescence method. Images were acquired on the Odyssey Fc imaging system (LI-COR) and band intensities were quantified using Image Studio Lite software (LI-COR).

### Primary neuronal cultures and transfection

Sprague-Dawley rat embryos (males and females, embryonic day 18) were used for the preparation of primary hippocampal and cortical neurons^60^. All research procedures involving the use of animals were conducted in accordance with the Australian Code of Practice for the Care and Use of Animals for Scientific Purposes and were approved by the University of Queensland Animal Ethics Committee (QBI/047/18). Briefly, hippocampi and cortices were isolated and dissociated separately with 30U of papain suspension (Worthington) for 20 min in a 37°C water bath. A single-cell suspension was obtained by triturating tissues with fire-polished glass Pasteur pipettes and then plated at a density of 8 × 10^4^ cells (for hippocampal neurons, per coverslip), 2 × 10^5^ cells (for cortical neurons, per well) or 1 × 10^6^ cells (for cortical neurons, per dish) on poly-L-lysine-coated 12-well plates or 6 cm dishes in Neurobasal growth medium supplemented with 2 mM Glutamax, 1% penicillin/streptomycin, and 2% B-27. Neurons were kept in a humidified 5% CO_2_ tissue culture incubator at 37°C. Hippocampal and cortical neurons were maintained in Neurobasal medium containing 0% or 1% FBS, respectively, and fed twice a week. For cortical neurons, uridine (Sigma, 5 μM) and 5’-fluoro-2’-deoxyuridine (Sigma, 5 μM) were added to the culture medium at days *in vitro* (DIV) 5 to stop glial proliferation. Transfection of hippocampal neurons was carried out at DIV 12-14 with Lipofectamine 2000 (Invitrogen) according to the manufacturer’s instructions. Cells were processed at DIV 15-17.

### Lentivirus Preparation and Transduction

HEK293T cells were transfected by the calcium-phosphate co-precipitation method using 10 μg of the plasmid of interest and 5 μg each of pMD2.G envelope plasmid, pRSV-Rev encoding plasmid and pMDLg/pRRE packaging construct^48,61^. Forty-eight hours after transfection, culture supernatants were collected and passed through a 0.45 μm cellulose acetate low binding protein membrane filter. Lentiviral particles were harvested either by ultracentrifugation at 106,559 *g* for 2 h at 4°C using a Beckman SW 32 Ti rotor or using the PEG-it Virus Precipitation Solution (System Biosciences) according to the manufacturer’s instructions. Concentrated virus particles were resuspended in Neurobasal medium, flash frozen with liquid nitrogen and stored at −80°C. Neurons were transduced with lentiviral particles between DIV 6-9 for 6 h or overnight, after which they were further incubated for 6-9 days prior to harvesting.

### Glycine-induced cLTP

cLTP was performed on primary rat hippocampal and cortical neurons at DIV 15-17 as previously described, with some modifications^61^. Briefly, neurons were washed and incubated in pre-warmed low Mg^2+^ artificial cerebrospinal fluid (ACSF; 120 mM NaCl, 2 mM CaCl_2_, 5 mM KCl, 0.4 mM MgCl_2_, 30 mM glucose, 25mM HEPES, pH 7.4) containing 0.5 µM tetrodotoxin (TTX, Tocris), 20 µM bicuculline (Abcam) and 5 µM strychnine (Sigma) at 37°C for 45 min. cLTP was induced by incubating neurons with low Mg^2+^ ACSF supplemented with 200 µM glycine, 20 µM bicuculline and 5 µM strychnine at room temperature for 5 min. To inhibit the activity of CaMKIIα, neurons were treated with KN-93 (Abcam, 10 μM) for 15 min prior to and during cLTP induction.

### Surface staining and antibody-feeding assays

To study the trafficking of NMDARs in primary hippocampal cultures, we use an antibody-feeding approach as described previously^62,63^. Surface GluN2A was labeled by incubating live primary hippocampal neurons transfected with various SEP-GluN2A constructs with rabbit anti-GFP antibody (JH4030, 1:250) for 30 min at 4°C prior to 10 min fixation in ice-cold parafix solution (4% paraformaldehyde, 4% sucrose in PBS). Following cell permeabilization (0.25% Triton X-100 in PBS, 10 min) and blocking (10% normal goat serum, 1 h) at room temperature, total SEP-GluN2A was labeled with chicken anti-GFP antibody (Abcam, 1:5,000) at 4°C overnight. The surface and total SEP-GluN2A were subsequently visualized by Alexa-568-conjugated anti-rabbit and Alexa-488-conjugated anti-chicken secondary antibodies, respectively. To determine the amount of receptor internalization, surface SEP-GluN2A was first labeled with rabbit anti-GFP antibody in live neurons at room temperature for 10 min. Neurons were washed with warm ACSF (25 mM HEPES, 120 mM NaCl, 5 mM KCl, 2 mM CaCl_2_, 2 mM MgCl_2_, 30 mM D-glucose, pH 7.4) and incubated at 37°C for 30 min to allow receptor internalization to occur. Surface-bound and non-internalized antibodies were blocked with an excess amount of unconjugated donkey anti-rabbit Fab fragments (Jackson ImmunoResearch, 1:50) at room temperature for 20 min. After washing, the neurons were fixed, blocked and incubated with Alexa-647-conjugated anti-rabbit secondary antibody at room temperature for 30 min. Neurons were subsequently washed, permeabilized, blocked and incubated with chicken anti-GFP antibody at 4°C overnight. The total and internalized SEP-GluN2A were visualized by Alexa-488-conjugated anti-chicken and Alexa-568-conjugated anti-rabbit secondary antibodies, respectively. To measure the level of receptor recycling, neurons expressing SEP-GluN2A were fed with rabbit anti-GFP antibody and allowed to be endocytosed for 30 min. After neurons were incubated with an excess amount of unconjugated donkey anti-rabbit Fab fragments, they were transferred back into a humidified 37°C tissue culture incubator for 30 min, which allowed internalized receptors to recycle back to the plasma membrane. Neurons were then fixed, blocked and incubated with Alexa-647-conjugated anti-rabbit secondary antibody which labels recycled receptors on the plasma membrane. They were subsequently washed, permeabilized, blocked and incubated with chicken anti-GFP antibody at 4°C overnight. The total and intracellular SEP-GluN2A were visualized by Alexa-488-conjugated anti-chicken and Alexa-568-conjugated anti-rabbit secondary antibodies, respectively. Images were collected with a 63X oil-immersion objective on a Zeiss LSM510 confocal microscope. Fluorescence intensities were quantified using Image J software (National Institutes of Health) for surface, internalized, and total GluN2A. Data were expressed as the surface/total GluN2A ratio, internalized/(surface+internalized) GluN2A (internalization index) or recycled/(surface+internalized) GluN2A (recycling index).

### Surface biotinylation assay

The surface biotinylation assay was carried out at DIV 15-17 to measure the amount of endogenous GluN2A protein on the plasma membrane^61^. Briefly, live neurons were washed once in ice-cold ACSF (pH 8.2), followed by a 30 min incubation with 1 mg/ml of sulfo-NHS-SS-biotin (CovaChem) at 4°C. Free biotin was quenched by washing the cells three times with ice-cold Tris-buffered saline (TBS, 50 mM Tris-HCl, 150 mM NaCl, pH 7.4). Neurons were lyzed in RIPA buffer (1% Triton X-100, 0.5% Na-deoxycholate, 0.1% SDS, 2 mM EDTA, 2 mM EGTA, 50 mM NaF, 10 mM Na-pyrophosphate in TBS, pH 7.4) supplemented with EDTA-free protease inhibitor cocktail (Sigma) at 4°C. Lysates were centrifuged at 20,627*g* for 20 min at 4°C and incubated with Neutravidin agarose beads (Thermo Scientific) overnight at 4°C. Beads were washed three times with ice-cold RIPA buffer and bound proteins were eluted with 2X SDS sample buffer at 50°C for 30 min and analyzed by western blotting. The absence of β-actin on the surface fraction confirmed the specificity of the assay.

### Immunoprecipitation assays

To determine the level of GluN2A-S1459 phosphorylation in primary cortical neurons, we performed immunoprecipitation assays using rabbit anti-GluN2A-pS1459 antibodies as previously described^28^. Briefly, neurons were lyzed in ice-cold RIPA buffer containing protease and phosphatase inhibitor cocktails. Lysates were centrifuged at 14,000 rpm for 20 min at 4°C and cleared with protein A-Sepharose beads (Thermo Scientific). Pre-cleared lysates were then incubated with anti-GluN2A-pS1459 coupled to protein A-Sepharose overnight at 4°C. After four washes with ice-cold lysis buffer, bound proteins were eluted in 2X SDS sample buffer at 50°C for 30 min. Both total and eluted proteins were analyzed by western blotting with specific antibodies against GluN2A, CaMKII and β-actin.

### Artificial synapse recordings

Methods for preparing neurons and HEK293 cells for artificial synapse recordings have previously been described^55^. Briefly, HEK293 cells were transfected with cDNAs encoding human GluN1 and SEP-tagged rat GluN2A subunits, plus empty pEGFP and mouse neuroligin-1B (Addgene #15261, in the pCAGGS expression vector) in a ratio of 1:1:1:0.5:1, using a calcium-phosphate co-precipitation protocol. Cortical neurons from embryonic day 18 rat embryos (ethics approval number: QBI/142/16/NHMRC/ARC) were plated on poly-D-lysine-coated coverslips at a density of 1 × 10^5^ cells per coverslip in DMEM supplemented with 10% FBS. The medium was replaced after 24 h with Neurobasal medium containing 2% B-27 and 1% Glutamax. After one week, half of this medium was replaced with fresh medium. Neurons were allowed to grow for 3-4 weeks before freshly transfected HEK293 cells were plated onto the neurons. Artificial synaptic connections typically formed spontaneously within 24 h and EPSCs in HEK293 cells were recorded 2-5 d later. Artificial synapse recordings were done in the whole-cell patch-clamp recording configuration at a holding potential of −70 mV unless otherwise stated, at 22 ± 2°C. Patch pipettes were fabricated from borosilicate hematocrit tubing (Harvard Apparatus) and had tip resistances of 2-5 MΩ when filled with the intracellular solution which contained (in mM): 145 CsCl, 2 CaCl_2_, 2 MgCl_2_, 10 HEPES, and 10 EGTA, adjusted to pH 7.4 with CsOH. Cells were perfused with extracellular solution, which contained (in mM): 140 NaCl, 5 KCl, 2 CaCl_2_, 10 HEPES and 10 D-glucose, adjusted to pH 7.4 with NaOH. EPSCs were filtered (−3dB, 4-pole Bessel) at 4 kHz, and then sampled at 10 kHz and recorded using a Multiclamp 700B amplifier and pClamp 10 software (Molecular Devices). Recordings with series resistances >20 MΩ were discarded and series resistance compensation was not applied to the recorded cell. Solutions containing defined concentrations of glutamate, glycine and Mg^2+^ were applied to cells by gravity-induced perfusion microtubules.

### Single-channel recordings

Single-receptor recordings were performed at room temperature (22 ± 1°C) in the outside-out patch configuration. The intracellular solution contained (in mM): 145 CsCl, 2 CaCl_2_, 2 MgCl_2_, 10 HEPES, and 10 EGTA, adjusted to pH 7.4 with CsOH. The extracellular solution comprised (in mM): 140 NaCl, 5 KCl, 2 CaCl_2_, 0.020 Na_2_EDTA, 10 HEPES, and 10 D-glucose. NaOH was used to adjust both solutions to a pH of 7.4. The concentration of EDTA was chosen to chelate Mg^2+^ and other contaminant divalent ions without affecting Ca^2+^. The single-channel conductance was determined by constructing current-voltage (I-V) plots from data obtained at clamped voltages of (in mV) ±70, ±35, ±15 and 0. The conductance was calculated using

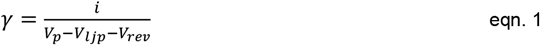

Where *V*_*p*_ is the holding potential (−70 mV), *V*_*ljp*_ is the liquid junction potential and *V*_*rev*_ is the reversal potential. *V*_*ljp*_ was calculated using JPCalWin and *V*_*rev*_ was estimated directly from the I-V plots. Single channel currents were recorded using an EPC 10 USB Heka Patch Clamp Amplifier, filtered (−3dB, 4-pole Bessel) at 2.9 kHz and sampled at 50 kHz. QuB software was used to determine the mean duration of single receptor activations and the intra-activation open probability (P_O_). Single channel currents were idealized using a resolution dead-time of 70 μs and the critical shut period that separated one discrete activation from the next was 200 ms. Activations with fewer than 20 events were excluded from analysis.

### Statistical Analysis

The sample size (*n*) reported in figure legends represents the number of individual neurons or wells generated from at least three independent experiments, unless otherwise stated. Statistical analysis was performed using Graph Pad Prism 8.0. One-way analysis of variance (ANOVA) with the Tukey’s, Sidak’s or Dunnet’s post-hoc multiple comparisons tests were used to compare the parameters for more than two groups. For the analyses of active duration and P_O_ of single-channel recordings, a Kruskal-Wallis one-way ANOVA with the Dunn’s post-hoc multiple comparisons test was used. For comparison between two groups, a two-tailed Mann-Whitney’s test was employed. All data are reported as mean ± standard error of the mean (SEM), or median with minimum and maximum values.

### Data availability

The data that support the findings of this study are available from the corresponding author upon reasonable request.

## Acknowledgements

We would like to thank Richard Huganir, David Baltimore, Peter Scheiffele, Salvatore Incontro and Patricio Opazo for providing DNA plasmids and antibodies, and Rowan Tweedale for critical reading of this manuscript. Imaging was performed at the Queensland Brain Institute’s Advanced Microscopy Facility. This work was supported by grants from the Australian Research Council (ARC, DP190101390, to V.A., B.M.C. and A.K.), the National Health and Medical Research Council (NHMRC, APP1099114, to B.M.C. and V.A.; and APP1156673 to A.K.), the National Institute of Neurological Disorders and Stroke Intramural Research Program (to K.W.R.), the Portuguese Foundation for Science and Technology (FCT [Fundação para a Ciência e a Tecnologia] grant SFRH/BI/106010/2015 to M.M.V.) and the Clem Jones Centre for Ageing Dementia Research to V.A. B.M.C. is supported by an NHMRC Senior Research Fellowship (APP1136021). J.W. is supported by a University of Queensland Amplify Fellowship and a previously held ARC Discovery Early Career Researcher Award (DE170100112). X.L.H.Y. is supported by a Research Training Program Scholarship from the Australian Government and the University of Queensland, as well as the Ian Lindenmayer PhD Top-up Scholarship.

## References

1. Nicoll RA, Roche KW. Long-term potentiation: peeling the onion. Neuropharmacology 74, 18–22 (2013).

2. Paoletti P, Bellone C, Zhou Q. NMDA receptor subunit diversity: impact on receptor properties, synaptic plasticity and disease. Nat Rev Neurosci 14, 383–400 (2013).

3. Morris RG. NMDA receptors and memory encoding. Neuropharmacology 74, 32–40 (2013).

4. Bosch M, Hayashi Y. Structural plasticity of dendritic spines. Curr Opin Neurobiol 22, 383–388 (2012).

5. Zhou Q, Sheng M. NMDA receptors in nervous system diseases. Neuropharmacology 74, 69–75 (2013).

6. XiangWei W, Jiang Y, Yuan H. De novo mutations and rare variants occurring in NMDA receptors. Curr Opin Physiol 2, 27–35 (2018).

7. Myers SJ, Yuan H, Kang JQ, Tan FCK, Traynelis SF, Low CM. Distinct roles of GRIN2A and GRIN2B variants in neurological conditions. F1000Res 8, (2019).

8. Stroebel D, Casado M, Paoletti P. Triheteromeric NMDA receptors: from structure to synaptic physiology. Curr Opin Physiol 2, 1–12 (2018).

9. Sanz-Clemente A, Nicoll RA, Roche KW. Diversity in NMDA receptor composition: many regulators, many consequences. Neuroscientist 19, 62–75 (2013).

10. Vieira M, Yong XLH, Roche KW, Anggono V. Regulation of NMDA glutamate receptor functions by the GluN2 subunits. J Neurochem 154, 121–143 (2020).

11. Wyllie DJ, Livesey MR, Hardingham GE. Influence of GluN2 subunit identity on NMDA receptor function. Neuropharmacology 74, 4–17 (2013).

12. Gray JA, Shi Y, Usui H, During MJ, Sakimura K, Nicoll RA. Distinct modes of AMPA receptor suppression at developing synapses by GluN2A and GluN2B: single-cell NMDA receptor subunit deletion in vivo. Neuron 71, 1085–1101 (2011).

13. Monyer H, Burnashev N, Laurie DJ, Sakmann B, Seeburg PH. Developmental and regional expression in the rat brain and functional properties of four NMDA receptors. Neuron 12, 529–540 (1994).

14. Sheng M, Cummings J, Roldan LA, Jan YN, Jan LY. Changing subunit composition of heteromeric NMDA receptors during development of rat cortex. Nature 368, 144–147 (1994).

15. Quinlan EM, Philpot BD, Huganir RL, Bear MF. Rapid, experience-dependent expression of synaptic NMDA receptors in visual cortex in vivo. Nat Neurosci 2, 352–357 (1999).

16. Bellone C, Nicoll RA. Rapid bidirectional switching of synaptic NMDA receptors. Neuron 55, 779–785 (2007).

17. Barria A, Malinow R. Subunit-specific NMDA receptor trafficking to synapses. Neuron 35, 345–353 (2002).

18. Grosshans DR, Clayton DA, Coultrap SJ, Browning MD. LTP leads to rapid surface expression of NMDA but not AMPA receptors in adult rat CA1. Nat Neurosci 5, 27–33 (2002).

19. Zhang XM, et al. Activity-induced synaptic delivery of the GluN2A-containing NMDA receptor is dependent on endoplasmic reticulum chaperone Bip and involved in fear memory. Cell Res 25, 818–836 (2015).

20. Swanger SA, He YA, Richter JD, Bassell GJ. Dendritic GluN2A synthesis mediates activity-induced NMDA receptor insertion. J Neurosci 33, 8898–8908 (2013).

21. Yashiro K, Philpot BD. Regulation of NMDA receptor subunit expression and its implications for LTD, LTP, and metaplasticity. Neuropharmacology 55, 1081–1094 (2008).

22. Kirkwood A, Rioult MC, Bear MF. Experience-dependent modification of synaptic plasticity in visual cortex. Nature 381, 526–528 (1996).

23. Barria A, Malinow R. NMDA receptor subunit composition controls synaptic plasticity by regulating binding to CaMKII. Neuron 48, 289–301 (2005).

24. Shipton OA, Paulsen O. GluN2A and GluN2B subunit-containing NMDA receptors in hippocampal plasticity. Philos Trans R Soc Lond B Biol Sci 369, 20130163 (2014).

25. Lussier MP, Sanz-Clemente A, Roche KW. Dynamic regulation of N-methyl-d-aspartate (NMDA) and α-amino-3-hydroxy-5-methyl-4-isoxazolepropionic acid (AMPA) receptors by posttranslational modifications. J Biol Chem 290, 28596–28603 (2015).

26. Cai L, Loo LS, Atlashkin V, Hanson BJ, Hong W. Deficiency of sorting nexin 27 (SNX27) leads to growth retardation and elevated levels of N-methyl-D-aspartate receptor 2C (NR2C). Mol Cell Biol 31, 1734–1747 (2011).

27. Clairfeuille T, et al. A molecular code for endosomal recycling of phosphorylated cargos by the SNX27-retromer complex. Nat Struct Mol Biol 23, 921–932 (2016).

28. Mota Vieira M, et al. An epilepsy-associated GRIN2A rare variant disrupts CaMKIIα phosphorylation of GluN2A and NMDA receptor trafficking. Cell Rep 32, 108104 (2020).

29. Gallon M, et al. A unique PDZ domain and arrestin-like fold interaction reveals mechanistic details of endocytic recycling by SNX27-retromer. Proc Natl Acad Sci U S A 111, E3604–3613 (2014).

30. Cullen PJ, Korswagen HC. Sorting nexins provide diversity for retromer-dependent trafficking events. Nat Cell Biol 14, 29–37 (2011).

31. Wang X, et al. Loss of sorting nexin 27 contributes to excitatory synaptic dysfunction by modulating glutamate receptor recycling in Down’s syndrome. Nat Med 19, 473–480 (2013).

32. Bowling KM, et al. Genomic diagnosis for children with intellectual disability and/or developmental delay. Genome Med 9, 43 (2017).

33. Lu W, Man H, Ju W, Trimble WS, MacDonald JF, Wang YT. Activation of synaptic NMDA receptors induces membrane insertion of new AMPA receptors and LTP in cultured hippocampal neurons. Neuron 29, 243–254 (2001).

34. Biederer T, Scheiffele P. Mixed-culture assays for analyzing neuronal synapse formation. Nat Protoc 2, 670–676 (2007).

35. Hussain NK, Diering GH, Sole J, Anggono V, Huganir RL. Sorting nexin 27 regulates basal and activity-dependent trafficking of AMPARs. Proc Natl Acad Sci U S A 111, 11840–11845 (2014).

36. Lauffer BE, et al. SNX27 mediates PDZ-directed sorting from endosomes to the plasma membrane. J Cell Biol 190, 565–574 (2010).

37. Traynelis SF, et al. Glutamate receptor ion channels: structure, regulation, and function. Pharmacol Rev 62, 405–496 (2010).

38. Wyllie DJ, Behe P, Colquhoun D. Single-channel activations and concentration jumps: comparison of recombinant NR1a/NR2A and NR1a/NR2D NMDA receptors. J Physiol 510 (Pt 1), 1–18 (1998).

39. Scott S, Lynch JW, Keramidas A. Correlating structural and energetic changes in glycine receptor activation. J Biol Chem 290, 5621–5634 (2015).

40. Dixon C, Sah P, Lynch JW, Keramidas A. GABAA receptor α and γ subunits shape synaptic currents via different mechanisms. J Biol Chem 289, 5399–5411 (2014).

41. Lavezzari G, McCallum J, Dewey CM, Roche KW. Subunit-specific regulation of NMDA receptor endocytosis. J Neurosci 24, 6383–6391 (2004).

42. Temkin P, Morishita W, Goswami D, Arendt K, Chen L, Malenka R. The retromer supports AMPA receptor trafficking during LTP. Neuron 94, 74–82 e75 (2017).

43. Loo LS, Tang N, Al-Haddawi M, Stewart Dawe G, Hong W. A role for sorting nexin 27 in AMPA receptor trafficking. Nat Commun 5, 3176 (2014).

44. Steinberg F, et al. A global analysis of SNX27-retromer assembly and cargo specificity reveals a function in glucose and metal ion transport. Nat Cell Biol 15, 461–471 (2013).

45. McMillan KJ, et al. Sorting nexin-27 regulates AMPA receptor trafficking through the synaptic adhesion protein LRFN2 bioRxiv, doi.org/10.1101/2020.04.27.063248 (2020).

46. Silva AJ, Stevens CF, Tonegawa S, Wang Y. Deficient hippocampal long-term potentiation in α-calcium-calmodulin kinase II mutant mice. Science 257, 201–206 (1992).

47. Bayer KU, Schulman H. CaM kinase: Still inspiring at 40. Neuron 103, 380–394 (2019).

48. Incontro S, et al. The CaMKII/NMDA receptor complex controls hippocampal synaptic transmission by kinase-dependent and independent mechanisms. Nat Commun 9, 2069 (2018).

49. Gardoni F, Bellone C, Cattabeni F, Di Luca M. Protein kinase C activation modulates α-calmodulin kinase II binding to NR2A subunit of N-methyl-D-aspartate receptor complex. J Biol Chem 276, 7609–7613 (2001).

50. Gardoni F, Schrama LH, van Dalen JJ, Gispen WH, Cattabeni F, Di Luca M. αCaMKII binding to the C-terminal tail of NMDA receptor subunit NR2A and its modulation by autophosphorylation. FEBS Lett 456, 394–398 (1999).

51. Kristensen AS, et al. Mechanism of Ca^2^+/calmodulin-dependent kinase II regulation of AMPA receptor gating. Nat Neurosci 14, 727–735 (2011).

52. Derkach V, Barria A, Soderling TR. Ca^2+^/calmodulin-kinase II enhances channel conductance of α-amino-3-hydroxy-5-methyl-4-isoxazolepropionate type glutamate receptors. Proc Natl Acad Sci U S A 96, 3269–3274 (1999).

53. Banke TG, Bowie D, Lee H, Huganir RL, Schousboe A, Traynelis SF. Control of GluR1 AMPA receptor function by cAMP-dependent protein kinase. J Neurosci 20, 89–102 (2000).

54. Li J, et al. De novo GRIN variants in NMDA receptor M2 channel pore-forming loop are associated with neurological diseases. Hum Mutat 40, 2393–2413 (2019).

55. Chen X, Keramidas A, Harvey RJ, Lynch JW. Effects of GluN2A and GluN2B gain-of-function epilepsy mutations on synaptic currents mediated by diheteromeric and triheteromeric NMDA receptors. Neurobiol Dis 140, 104850 (2020).

56. Chih B, Gollan L, Scheiffele P. Alternative splicing controls selective trans-synaptic interactions of the neuroligin-neurexin complex. Neuron 51, 171–178 (2006).

57. Opazo P, et al. CaMKII triggers the diffusional trapping of surface AMPARs through phosphorylation of stargazin. Neuron 67, 239–252 (2010).

58. Anggono V, Clem RL, Huganir RL. PICK1 loss of function occludes homeostatic synaptic scaling. J Neurosci 31, 2188–2196 (2011).

59. Yong XLH, Cousin MA, Anggono V. PICK1 controls activity-dependent synaptic vesicle cargo retrieval. Cell Rep 33, 108312 (2020).

60. Widagdo J, et al. Activity-dependent ubiquitination of GluA1 and GluA2 regulates AMPA receptor intracellular sorting and degradation. Cell Rep 10, 783–795 (2015).

61. Tan MC, et al. The activity-induced long non-coding RNA Meg3 modulates AMPA receptor surface expression in primary cortical neurons. Front Cell Neurosci 11, 124 (2017).

62. Chiu AM, Barse L, Hubalkova P, Sanz-Clemente A. An antibody feeding approach to study glutamate receptor trafficking in dissociated primary hippocampal cultures. J Vis Exp 150, e59982 (2019).

63. Anggono V, et al. PICK1 interacts with PACSIN to regulate AMPA receptor internalization and cerebellar long-term depression. Proc Natl Acad Sci U S A 110, 13976–13981 (2013).

