## Supplementary Figures and Tables for "Regulation of NMDA receptor trafficking and gating by activity-dependent CaMKIIα phosphorylation of the GluN2A subunit"

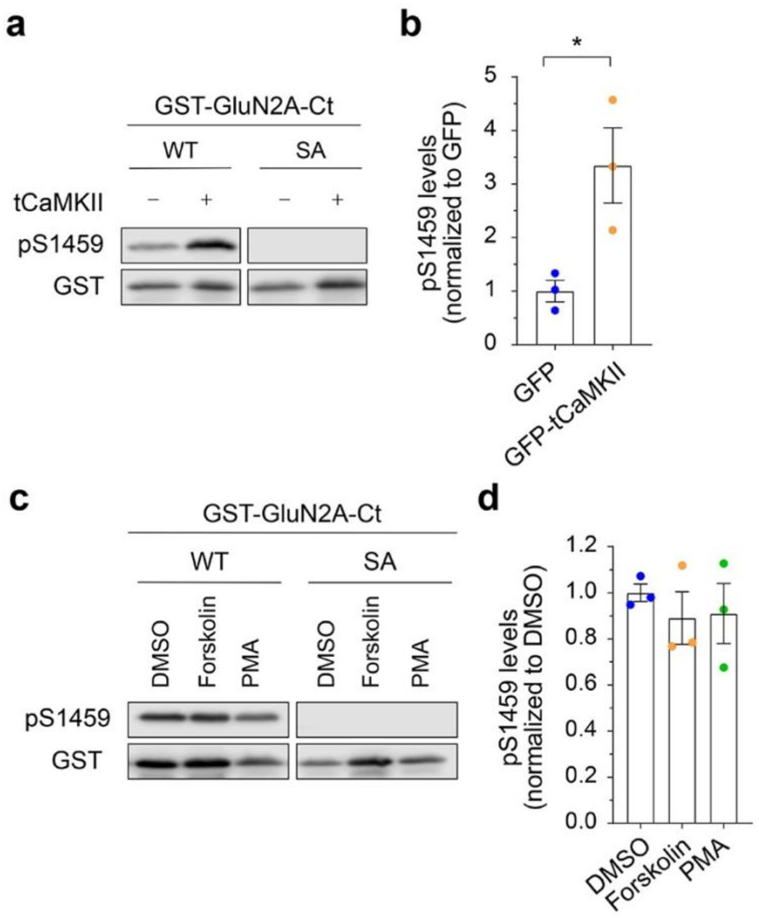


**Supplementary Fig. 1.** CaMKIIα phosphorylates GluN2A on Ser-1459. **a** HEK293T cells were transfected with pCIS-GluN2A-C-terminal domain (WT or S1459A) and pEGFP or pEGFP-tCaMKIIα (constitutively active truncated CaMKIIα), lyzed and pulled-down with GSH-sepharose. Bound proteins were eluted and resolved by SDS-PAGE, and analyzed by western blotting with specific antibodies against GluN2A pS1459 and GST. **b** Quantification of Ser-1459 phosphorylation levels after normalizing to GST. Data represent mean ± SEM of band intensities normalized to control values of cells expressing GFP ($n$ = 3). *$P$ < 0.05 using unpaired $t$-test. **c** HEK293T cells expressing GST-GluN2A-C-terminal domain (WT or S1459A) were treated with 10 µM forskolin, 1 µM PMA or DMSO (vehicle) for 10 min prior to lysis. Cell lysates were analyzed by immunoblotting with specific antibodies against GluN2A pS1459 and GST. **d** Quantification of Ser-1459 phosphorylation levels after normalizing to GST. Data represent mean ± SEM of band intensities normalized to control values of vehicle-treated cells ($n$ = 3). No statistical difference was observed among groups as determined by one-way ANOVA.

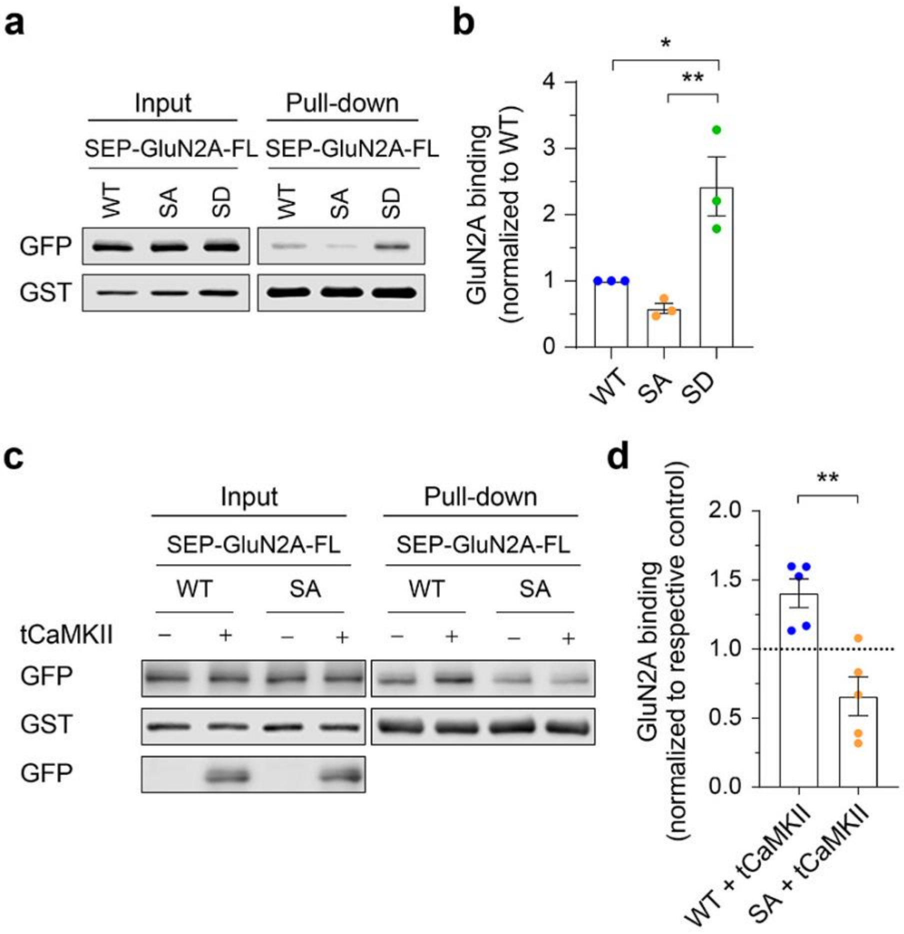


**Supplementary Fig. S2.** GluN2A phosphorylation on Ser-1459 enhances SNX27 binding. **a** HEK293T cells were co-transfected with plasmids encoding GST-SNX27, GluN1 and full-length SEP-GluN2A (WT, S1459A or S1459D), lyzed and pulled-down with GSH-sepharose. Bound proteins were eluted and resolved by SDS-PAGE, and analyzed by western blotting with specific antibodies against GFP and GST. **b** Quantification of the level of SEP-GluN2A binding to GST-SNX27. Data represent mean ± SEM of band intensities normalized to control values of cells expressing SEP-GluN2A WT ($n$ = 3). *$P < 0.05$, **$P < 0.01$ using one-way ANOVA with Tukey's multiple comparisons test. **c** HEK293T cells were co-transfected with plasmids encoding GST-SNX27, GluN1 and SEP-GluN2A (WT or S1459A), with pEGFP or pEGFP-tCaMKIIα. Cells were lyzed and pulled-down with GSH-sepharose. Bound proteins and total lysates were analyzed by immunoblotting with specific antibodies against GFP and GST. **d** Quantification of the level of SEP-GluN2A binding to GST-SNX27. Data represent mean ± SEM of band intensities normalized to the respective control values of cells that expressed GFP only ($n$ = 5). **$P < 0.01$ using Mann-Whitney test.

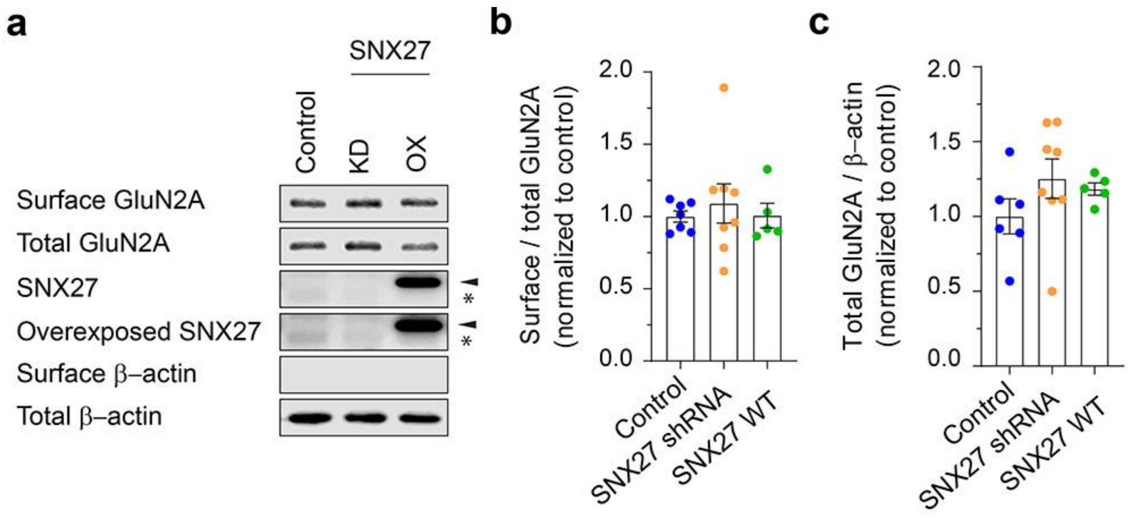


**Supplementary Fig. 3.** SNX27 does not regulate GluN2A surface expression under basal conditions. **a** Cultured cortical neurons were transduced with lentiviral particles expressing GFP (control), SNX27 shRNA or myc-SNX27 cDNA (OX) at DIV 9. At DIV15, neurons were subjected to surface biotinylation assays. The relative amount of surface and total GluN2A was assessed by western blotting using specific antibodies against GluN2A. SNX27 blots confirmed the levels of knockdown and overexpression. Endogenous and overexpressed SNX27 are indicated by asterisks and arrowheads, respectively. Quantification of the surface/total ratio (*B*) and total (*C*) GluN2A after normalizing against β-actin. Data represent mean ± SEM of band intensities normalized to control values of neurons expressing GFP only ($n$ = 5-8). No statistical difference was observed among groups as determined by one-way ANOVA.

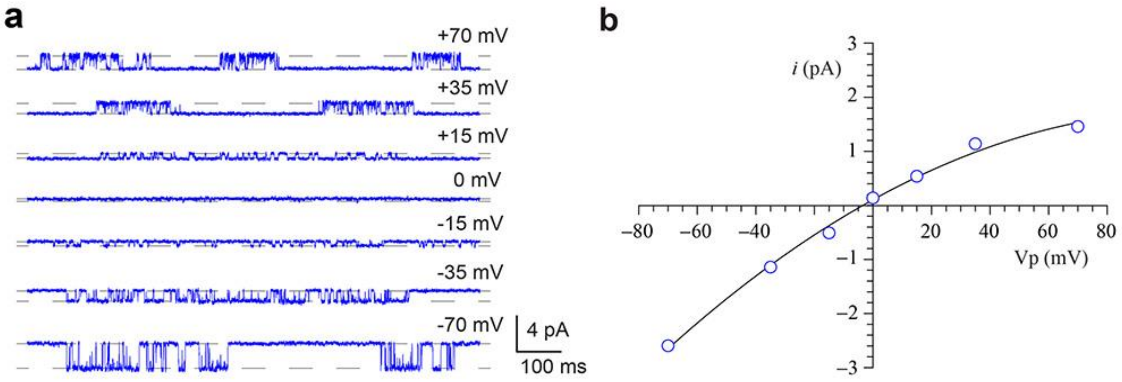


**Supplementary Fig. 4.** Single channel I-V relationships. **a** Single channel current amplitude as a function of voltage mediated by wild-type GluN1/2A receptors from a single excised patch of membrane. Single receptor currents were elicited by continuously perfusing the recorded patch with 1 mM glutamate, 0 $Mg^{2+}$ and 100 μM glycine at the indicated applied voltages. **b** The I-V curve.

**Supplementary Fig. 5**. Uncropped images of western blots.

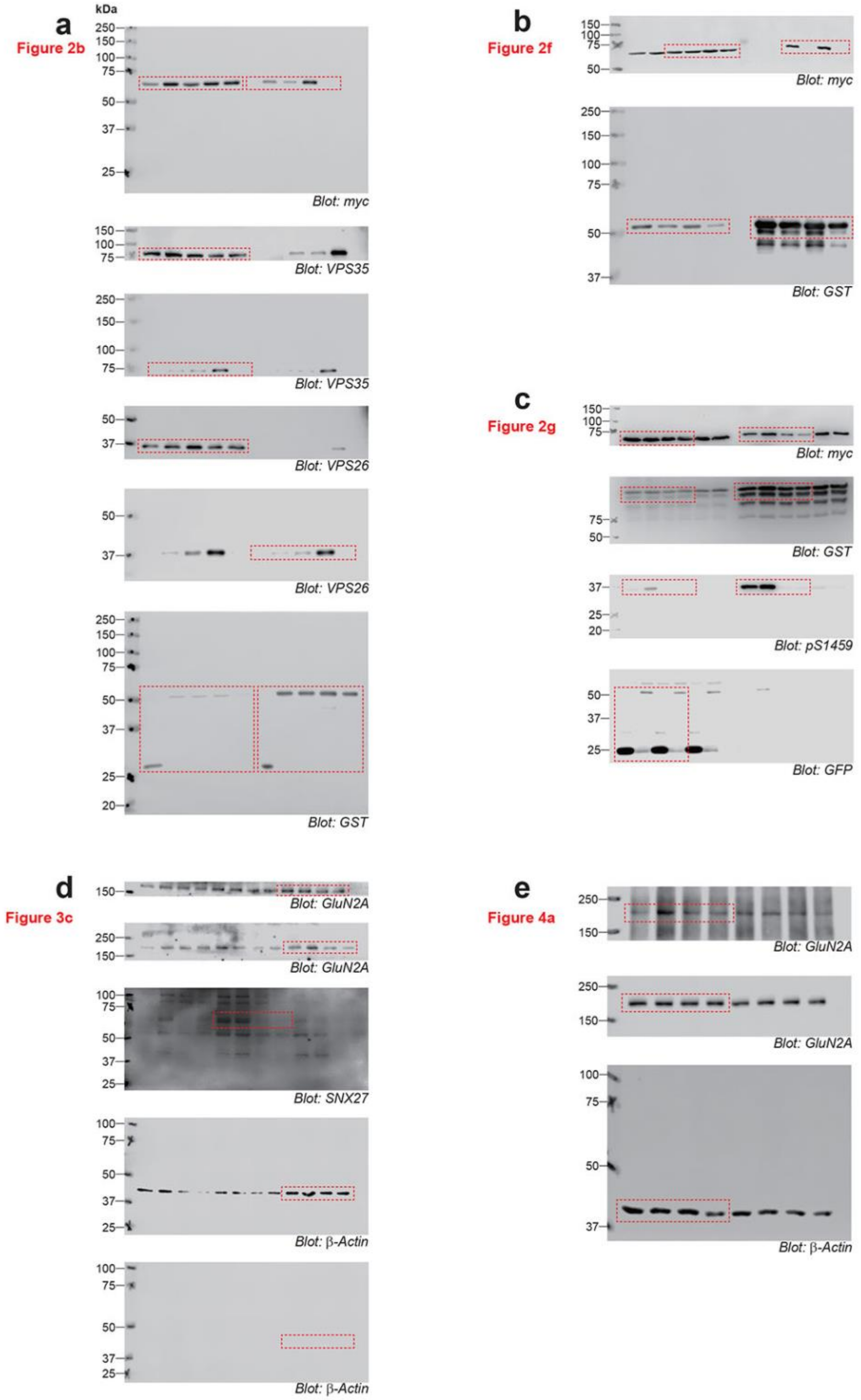

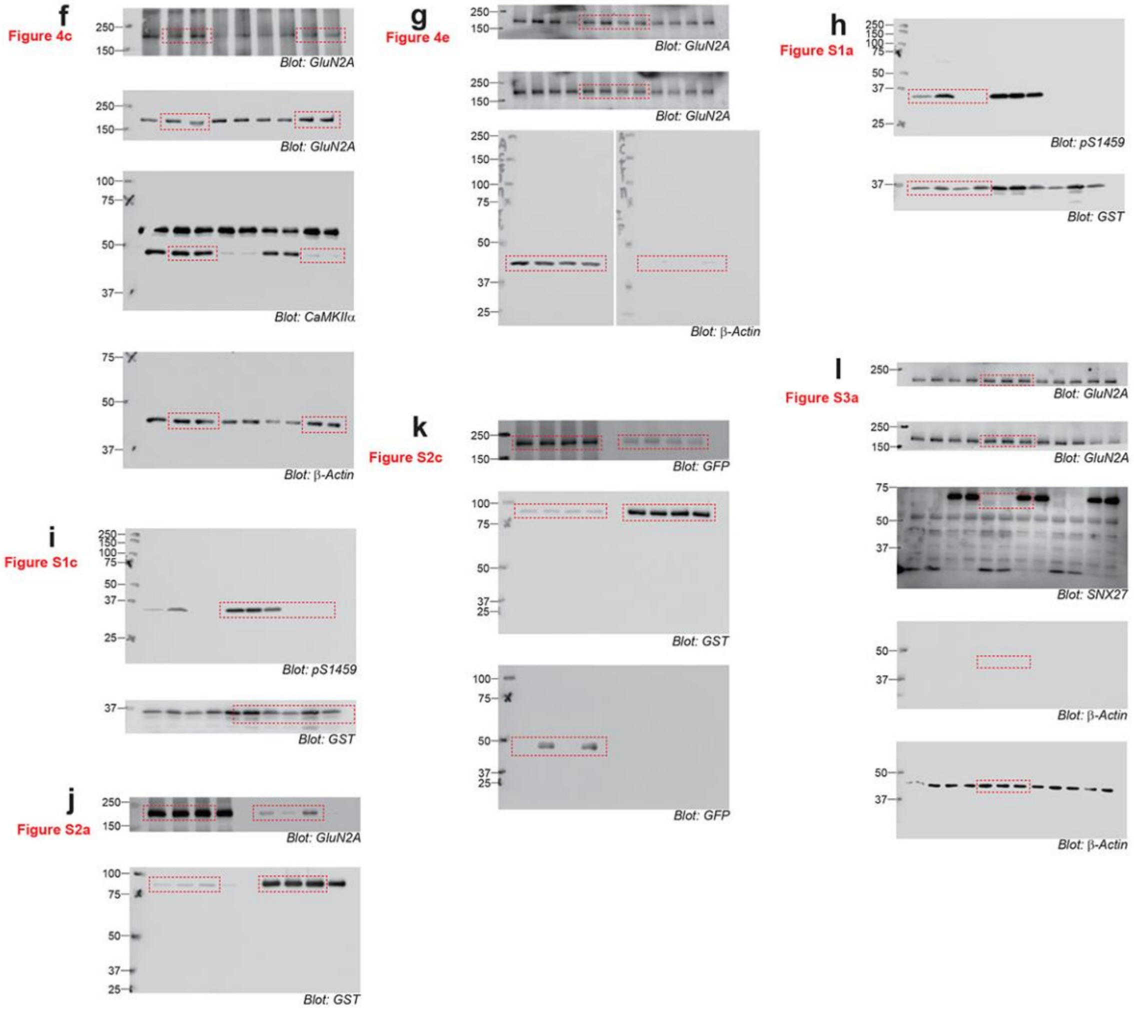
f
Figure 4c
Blot: GluN2A
Blot: GluN2A
Blot: CaMKIIα
Blot: β-Actin
g
Figure 4e
Blot: GluN2A
Blot: GluN2A
Blot: β-Actin
h
Figure S1a
Blot: pS1459
Blot: GST
i
Figure S1c
Blot: pS1459
Blot: GST
j
Figure S2a
Blot: GluN2A
Blot: GST
k
Figure S2c
Blot: GFP
Blot: GST
Blot: GFP
l
Figure S3a
Blot: GluN2A
Blot: GluN2A
Blot: SNX27
Blot: β-Actin
Blot: β-Actin

**Supplementary Table 1.** ITC binding data for SNX27 association with GluN2A C-terminal peptides.

| Peptide | Sequence | Kd, μM | Δ*H*, kcal/mol | TΔ*S*, kcal/mol | Δ*G*, kcal/mol | N, stoichiometry |
|---|---|---|---|---|---|---|
| WT | YKKMPSIESDV | 46.0 ± 2.6 | -5.8 ± 1.9 | -0.13 ± 0.02 | -5.9 ± 1.0 | 1.2 ± 0.1 |
| S1459A | YKKMPAIESDV | 49.9 ± 1.3 | -5.3 ± 1.2 | 0.53 ± 0.03 | -5.9 ± 0.9 | 1.1 ± 0.1 |
| S1459E | YKKMPEIESDV | 25.0 ± 2.1 | -17.1 ± 1.6 | -10.88 ± 1.02 | -6.2 ± 0.4 | 0.9 ± 0.1 |
| S1462E | YKKMPSIEEDV | n.b.d. | n.b.d. | n.b.d. | n.b.d. | n.b.d. |
| p-S1459 | YKKMP(pS)IESDV | 13.7 ± 0.7 | -9.0 ± 1.0 | -2.2 ± 0.9 | -6.6 ± 0.03 | 0.7 ± 0.1 |

All experiments were performed 3 times. Data are presented as mean ± SD. n.b.d. = no binding detectable.

**Supplementary Table 2.** Single channel conductance at -70 mV.

| Receptors | Conductance, pS |
|---|---|
| GluN1/GluN2A$^{WT}$ | 41.5 |
| GluN1/GluN2A$^{S1459G}$ | 42.7 |
| GluN1/GluN2A$^{S1459D}$ | 42.7 |
